# HSF1 inhibition potentiates the antifungal activity and proteome instability induced by the fungal Hsp110 inhibitor 2H

**DOI:** 10.64898/2026.10.12.758498

**Authors:** Cancan Sun, Balaji Nagarajan, Arjun Rijal, Charles Lyons, Abu Hamza, Wei Wang, Xianjun Fang, Yana Cen, Qinglian Liu

## Abstract

Fungal infections remain a major cause of morbidity and mortality worldwide; however, therapeutic options are limited and are increasingly compromised by antifungal resistance. Heat Shock Protein 110 kDa (Hsp110) is an essential molecular chaperone required for protein homeostasis (proteostasis) and fungal viability and has recently emerged as a promising antifungal target. We previously identified 2H, a first-in-class inhibitor of fungal Hsp110 proteins that exhibits broad-spectrum antifungal activity through disruption of proteostasis. Heat Shock Transcription Factor 1 (HSF1), the master regulator of cellular stress responses and chaperone expression, has also been proposed as a potential antifungal target, although direct pharmacological evidence supporting this strategy remains limited. Here, we investigated the effects of combining the HSF1 inhibitor DTHIB with the fungal Hsp110 inhibitor 2H in *Cryptococcus neoformans* and *Candida albicans*. Although DTHIB alone exhibited minimal antifungal activity, it significantly enhanced the growth-inhibitory and fungicidal effects of 2H. Notably, this potentiating effect was specific to 2H and was not observed with fluconazole. Consistent with the roles of Hsp110 and HSF1 in proteostasis, 2H induced extensive accumulation of detergent-insoluble protein aggregates in *C. neoformans*, whereas HSF1 inhibition further increased both the abundance and diversity of aggregated proteins. Quantitative proteomic analyses demonstrated that 2H induces proteome-wide protein aggregation, impacting proteins involved in a broad range of essential cellular processes, with high-molecular-weight proteins exhibiting particular susceptibility. Together, these findings provide pharmacological evidence supporting HSF1 as a promising antifungal sensitization target and demonstrate the utility of 2H as a chemical probe for studying Hsp110-dependent proteostasis. Simultaneous disruption of both HSF1-mediated stress responses and Hsp110-dependent proteostasis promotes extensive proteome instability and enhances antifungal activity, highlighting a potential new strategy for antifungal drug development.

**Author summary:** Fungal infections caused by pathogens such as *Cryptococcus neoformans* and *Candida albicans* are a major cause of illness and death worldwide, and treatment options are increasingly limited by antifungal resistance. New therapeutic strategies are urgently needed. We previously identified 2H, a first-in-class inhibitor of fungal Hsp110 proteins, molecular chaperones that help maintain protein homeostasis. In this study, we investigated whether inhibition of Heat Shock Transcription Factor 1 (HSF1), a key regulator of cellular stress responses and chaperone expression, could enhance the antifungal activity of Hsp110 inhibition. We found that the HSF1 inhibitor DTHIB significantly increased the antifungal and fungicidal activities of 2H and enhanced the protein aggregation caused by Hsp110 inhibition. Quantitative proteomic analyses revealed widespread disruption of protein homeostasis affecting multiple essential cellular pathways. Our findings show that fungal cells rely on HSF1-mediated stress responses to withstand the proteotoxic stress caused by Hsp110 inhibition. Simultaneous disruption of both pathways triggers extensive proteome instability and enhances antifungal activity, suggesting a new strategy for antifungal drug development.

## INTRODUCTION

Fungal diseases are an increasingly urgent global health challenge, causing an estimated 3.8 million deaths annually worldwide, with mortality nearly doubling over the past decade(1). Invasive fungal infections remain a leading cause of morbidity and mortality among immunocompromised individuals, including patients with HIV/AIDS, cancer, organ transplantation, or immunosuppressive therapies (2–7). Among the most clinically important pathogens are *Candida* species and *Cryptococcus neoformans* (6, 8–12), both recognized by the World Health Organization as priority fungal pathogens because of their substantial disease burden and limited treatment options. Despite their impact on human health, fungal diseases remain relatively understudied compared with bacterial and viral infections.

Current antifungal therapy relies primarily on azoles, polyenes, echinocandins, and flucytosine (9, 13–18). These agents target only a small number of fungal pathways, including ergosterol biosynthesis, cell wall assembly, and nucleic acid synthesis. The growing prevalence of antifungal resistance, together with the toxicity and pharmacological limitations of existing therapies, has reduced treatment options. Furthermore, the close evolutionary relationship between fungi and humans limits the availability of pathogen-specific drug targets. Consequently, there is an urgent need to develop antifungal agents with novel mechanisms of action.

Protein homeostasis (proteostasis) is essential for cellular viability and adaptation to environmental stress (19–24). Because pathogenic fungi encounter substantial proteotoxic stress during infection, molecular chaperones and protein quality-control pathways have emerged as attractive targets for antifungal intervention (25–27). Heat Shock Proteins 110 kDa (Hsp110s) are conserved molecular chaperones in eukaryotes that participate in diverse processes required for proteostasis, including *de novo* protein folding and stress-induced refolding, protein import into the endoplasmic reticulum (ER), protein degradation, and the prevention and solubilization of protein aggregates (28–53). The sole Hsp110 homolog in *Candida albicans*, Msi3, is essential for fungal growth, viability, and virulence, highlighting the importance of Hsp110-dependent proteostasis in fungal biology (28, 54–56).

Ubiquitously present in the cytosol of eukaryotes, Hsp110s function both as independent chaperones and as co-chaperones for Hsp70s (29, 31, 42, 44, 57). As independent chaperones, Hsp110s exhibit potent chaperone activity in preventing the aggregation of denatured proteins (36, 40, 50, 58, 59). As co-chaperones, Hsp110s serve as the major nucleotide-exchange factors (NEFs) for Hsp70s (42, 44, 60–62). While the molecular mechanism and biological importance of the NEF activity of Hsp110s are well established (63, 64), the role of their chaperone activity remains largely unclear. Despite the central role of Hsp110s in maintaining proteome integrity, the endogenous proteins that depend on their chaperone activity remain poorly defined. Identification of Hsp110-dependent proteins has proven difficult because interactions between Hsp110s and their client proteins are thought to be highly dynamic and transient (28, 65). As a result, how Hsp110s contribute to proteome homeostasis *in vivo* remains incompletely understood.

We recently identified 2H, a pyrazolo[3,4-b]pyridine derivative, as a first-in-class inhibitor of fungal Hsp110s. 2H selectively inhibits the chaperone activity of Msi3, the sole Hsp110 in *C. albicans*, while largely sparing its NEF activity. Importantly, 2H exhibits broad-spectrum antifungal activity against multiple pathogenic fungi (66, 67). Mechanistically, 2H disrupts proteostasis and induces extensive protein aggregation. Beyond its therapeutic potential, 2H provides a unique chemical tool for investigating the chaperone activity of Hsp110s. Proteins that lose solubility following Hsp110 inhibition may depend directly or indirectly on Hsp110-mediated proteostasis. Therefore, aggregate proteomics offers a powerful strategy for identifying candidate Hsp110-dependent proteins and elucidating the biological functions of Hsp110s *in vivo*.

Heat Shock Transcription Factor 1 (HSF1) is the master transcriptional regulator of the heat-shock response and coordinates adaptive responses to proteotoxic stress (68–79). HSF1 induces the expression of molecular chaperones and other protein quality-control factors, including members of the Hsp70, Hsp90, and Hsp110 families, thereby increasing cellular capacity to maintain proteome integrity. In fungal pathogens, HSF1 is essential for growth, stress adaptation, and virulence (80–82), whereas mammalian cells remain viable in its absence under normal conditions (71, 83, 84), making it an attractive candidate target for antifungal development. However, despite this compelling biological rationale, direct evidence supporting pharmacological inhibition of HSF1 as an antifungal strategy remains limited. The lack of effective approaches for selectively inhibiting fungal HSF1 has made it difficult to evaluate its therapeutic potential. Consequently, whether HSF1 inhibition can be exploited to enhance antifungal efficacy has remained largely unresolved.

Because HSF1 functions upstream of the cellular chaperone machinery, we hypothesized that inhibition of HSF1 would compromise adaptive proteostasis responses and sensitize fungal cells to Hsp110 inhibition. In addition, treatment with 2H induces compensatory Hsp110 expression in response to Hsp110 inhibition(66). DTHIB (Direct Targeted HSF1 InhiBitor) is a small-molecule inhibitor originally developed against human HSF1 (85). Although originally developed as an anticancer agent, its ability to suppress HSF1-dependent stress responses makes it a valuable pharmacological tool for probing HSF1 function in fungal cells.

In this study, we investigated the effects of simultaneously targeting HSF1 and Hsp110 in *C. neoformans*. We demonstrate that DTHIB potentiates the antifungal and fungicidal activities of the Hsp110 inhibitor 2H and significantly enhances protein aggregation induced by Hsp110 inhibition. In contrast, no enhancement of antifungal activity was observed with fluconazole, the most common antifungal medication, supporting the specificity of this enhancement. Quantitative proteomic analyses revealed widespread disruption of protein solubility across various essential cellular pathways and cellular compartments, particularly proteins with high molecular weights. These findings suggest that fungal cells rely on HSF1-mediated stress responses to withstand the proteotoxic stress caused by Hsp110 inhibition. In addition, the aggregate proteome generated following Hsp110 inhibition provides an important resource for identifying Hsp110-dependent proteins and elucidating the biological functions of Hsp110s *in vivo*. Together, these results support dual targeting of HSF1 and Hsp110 as a promising antifungal strategy and establish a framework for future mechanistic studies of Hsp110 biology. Importantly, this approach targets a fundamental biological process distinct from those affected by existing antifungals, which primarily target the cell wall, cell membrane, or nucleic acid metabolism (9, 13–15).

## RESULTS

### DTHIB potentiates the antifungal activity of the Hsp110 inhibitor 2H against Cryptococcus neoformans

To determine whether inhibition of HSF1 enhances the antifungal activity of Hsp110 inhibition, we evaluated the effects of combining the HSF1 inhibitor DTHIB with the fungal Hsp110 inhibitor 2H. The chemical structures of 2H and DTHIB are shown in **Fig. 1A**. Previous studies have shown that DTHIB binds to the DNA-binding domain (DBD) of human HSF1 (hHSF1) and promotes HSF1 degradation while having little impact on DNA binding (85). The DBDs of HSF1 proteins from *C. neoformans* (cnHSF1) and *C. albicans* (CTA8) share substantial sequence conservation with hHSF1, particularly cnHSF1 (**Fig. S1**), suggesting that DTHIB may interact with these fungal HSF1 in a similar manner.

**Figure 1.**
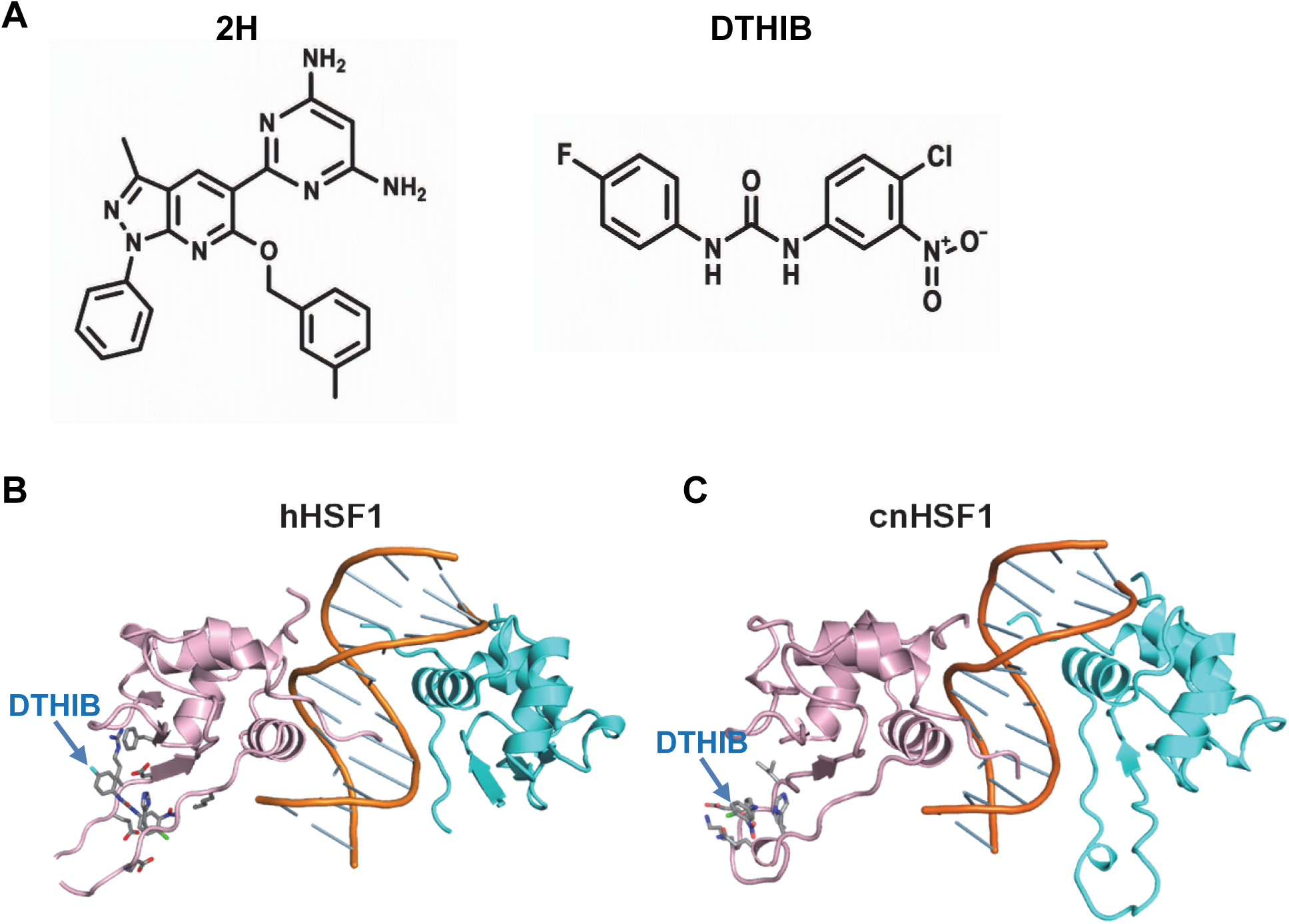
DTHIB is predicted to bind fungal HSF1 in a manner similar to that of human HSF1. **(A)** Chemical structures of 2H (left) and DTHIB (right). **(B,C)** Predicted binding modes of DTHIB in human HSF1 (hHSF1) **(B)** and *C. neoformans* HSF1 (cnHSF1) **(C)**. Molecular docking was performed using the crystal structure of the hHSF1-DNA complex (PDB: 5D5V) and an AlphaFold-predicted structure of cnHSF1 aligned to the hHSF1-DNA complex. Each structure contains two HSF1 protomers (shown in pink and cyan). DTHIB was docked to the pink protomer, which is more completely resolved in the hHSF1-DNA structure. The DNA backbone is shown in gold. DTHIB and interacting residues are displayed as sticks. The hHSF1 and cnHSF1 structures were superimposed using the pink protomer for structural alignment. Detailed protein-ligand interactions are shown in **Fig. S2**.

Since the precise DTHIB-binding site on hHSF1 has not been experimentally defined, we carried out *in silico* docking analyses to predict its binding mode using a published DNA-bound hHSF1 structure (PDB code: 5D5V(86)). Docking predicted that DTHIB binds to a surface-exposed loop region located distal to the DNA-binding interface (**Fig. 1B**). Thus, this binding mode is consistent with previous reports that DTHIB does not interfere with DNA binding.

As no experimental structure is available for cnHSF1, we generated an AlphaFold model and positioned two cnHSF1 DBDs onto the DNA duplex using the hHSF1-DNA complex as a template. Docking analysis predicted a similar DTHIB-binding mode for cnHSF1 (**Fig. 1C**). The predicted binding sites in hHSF1 and cnHSF1 exhibited similar AutoDock Vina scores (-5.65 and -5.98 kcal/mol, respectively). Although two interacting residue pairs, H83/H148 and E85/Q73, were conserved between hHSF1 and cnHSF1, the identities of the remaining interacting residues were largely divergent (**Fig. S2**). Nevertheless, the predicted binding environments exhibited similar physicochemical properties, including hydrophobic, polar, acidic, and basic features (**Fig. S2**). PLIP analysis further indicated that DTHIB occupies structurally analogous pockets in both proteins. Together, these findings suggest that DTHIB may inhibit cnHSF1 through a mechanism similar to that proposed for hHSF1.

Based on the high degree of conservation between hHsf1 and cnHSF1 (**Fig. S1, 1B-C**), we next examined the activity of DTHIB against *C. neoformans*. *C. neoformans* is a major cause of fungal meningitis and disseminated cryptococcosis, particularly in immunocompromised patients (11, 12, 87, 88). The widely used laboratory strain H99 was tested. DTHIB alone had little effect on fungal growth at the concentrations examined (**Fig. 2A**), suggesting that inhibition of HSF1-dependent stress responses is insufficient to substantially impair growth under these conditions. In contrast, DTHIB significantly enhanced the activity of 2H, resulting in a pronounced leftward shift of the 2H dose-response curve (**Fig. 2A**). Two-way ANOVA revealed significant effects of 2H concentration (F(7,120)=298.7, P<0.0001), DTHIB treatment (F(2,120)=121.9, P<0.0001), and a significant interaction between the two factors (F(14,120)=23.37, P<0.0001). Post hoc Sidak analysis demonstrated significant enhancement of 2H activity by DTHIB at concentrations between 1.56 and 12.5 µM (**Table S1**). Bliss independence analysis demonstrated concentration-dependent synergistic interactions between 2H and DTHIB. Synergy was more pronounced in the presence of 50 μM DTHIB (**Fig. 2B, Table S3**), whereas 25 μM DTHIB produced significant positive Bliss scores only at 6.25 and 12.5 μM (**Fig. S3A; Table S4**), indicating weaker but detectable synergistic interactions.

**Figure 2.**
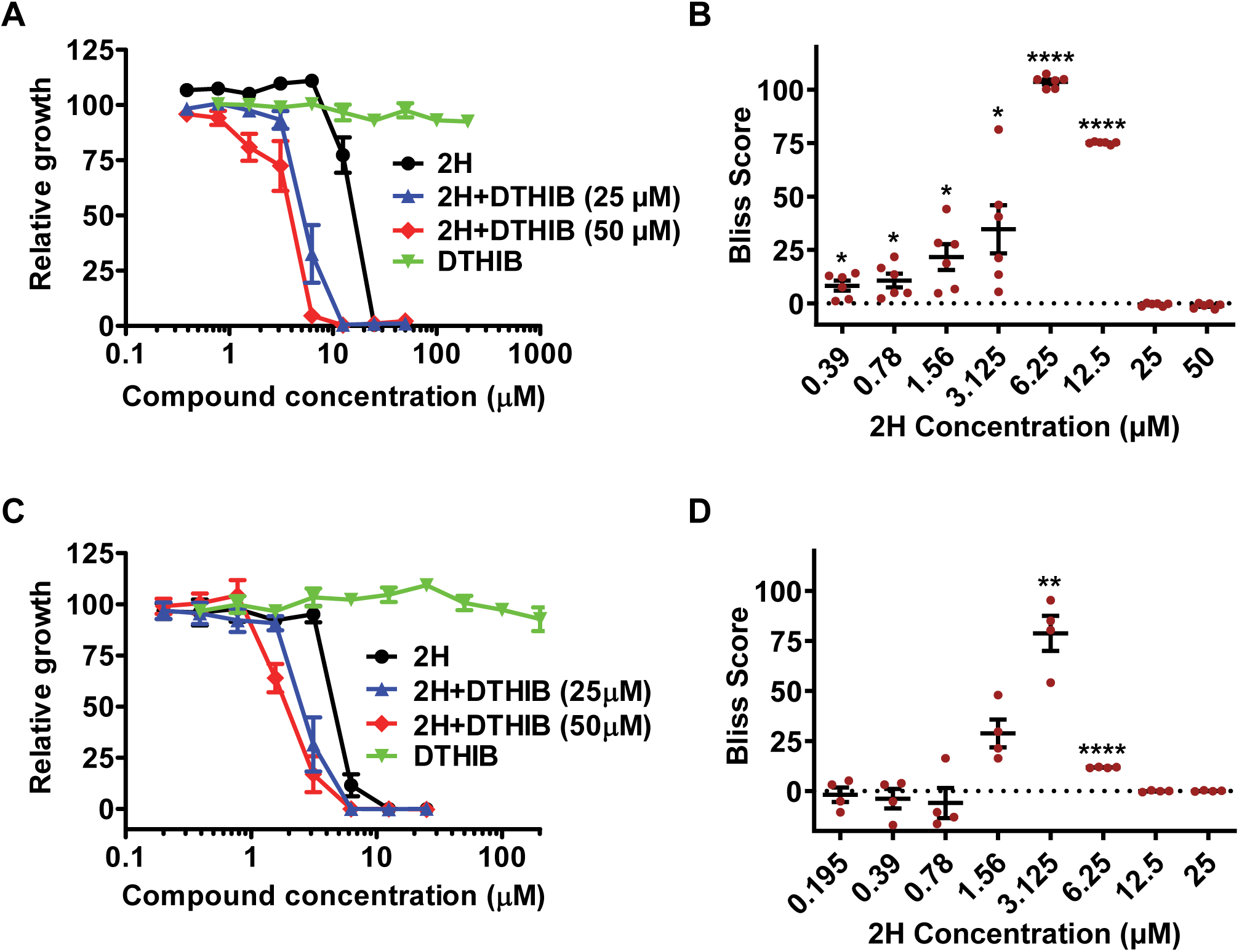
DTHIB potentiates the antifungal activity of the Hsp110 inhibitor 2H against both *C. neoformans* and *C. albicans*. **(A,C)** Growth inhibition of *C. neoformans* strain H99 **(A)** and *C. albicans* strain SC5314 **(C)** by 2H in the presence or absence of DTHIB. DTHIB alone exhibited minimal effects on fungal growth but enhanced the antifungal activity of 2H. Relative growth was normalized to the vehicle-treated control (100%). Data are presented as mean ± SEM from independent biological replicates (n=6 for *C. neoformans* and n=4 for *C. albicans*). Statistical significance was assessed by ordinary two-way ANOVA followed by Sidak’s multiple-comparison test. Detailed pairwise comparisons are provided in **Table S1** and **S2**. **(B,D)** Bliss synergy analysis of the data shown in **(A)** and **(C)**, respectively, using 50 μM DTHIB. Bliss scores were calculated using the Bliss independence model as expected relative growth minus observed relative growth. Positive, zero, and negative Bliss scores indicate synergistic, additive, and antagonistic interactions, respectively. The dashed line indicates Bliss = 0. Data are presented as mean ± SEM (n = 6 for *C. neoformans* and n = 4 for *C. albicans*). Bliss scores were compared with the theoretical value of zero, and P values were adjusted using the Benjamini-Hochberg false discovery rate (FDR) procedure. *P < 0.05; **P < 0.01; ****P < 0.0001. Detailed statistical results are provided in **Tables S3** and **S5**. Bliss analyses using 25 μM DTHIB are shown in **Figs. S3A** and **S3B**, with corresponding statistics in **Tables S4** and **S6**.

Consistent with the growth inhibition data, DTHIB lowered both the minimum inhibitory concentration for 90% inhibition (MIC_90_) and minimum fungicidal concentration (MFC) of 2H (**Fig. 2A**, **Table 1**). Under the conditions used in this study, 2H alone exhibited an MIC_90_ and MFC of 25 μM. Addition of 25 μM DTHIB reduced both values by approximately two-fold, whereas 50 μM DTHIB reduced the MIC_90_ to 6.25 μM and the MFC to 12.5 μM. Thus, inhibition of HSF1 significantly increased the susceptibility of *C. neoformans* to Hsp110 inhibition.

**Table 1.** DTHIB specifically enhances the antifungal activity of the Hsp110 inhibitor 2H, but not fluconazole. MIC_90_ and MFC values for 2H alone or in combination with DTHIB are listed. Fluconazole (FLC) was used as a control. MIC_90_: the minimum inhibitory concentration for 90% inhibition. The minimal fungicidal concentration (MFC): the minimal concentration of drug resulting in less than 0.1% of the cells alive relative to the original inoculum (i.e., >99.9% reduction relative to the starting inoculum). NA: not applicable due to the fungistatic nature of fluconazole.

| Strains | Compounds | MIC<sub>90</sub> (μM) | MFC (μM) |
| --- | --- | --- | --- |
| *C. neoformans* | 2H | 25 | 25 |
|  | 2H+25 μM DTHIB | 12.5 | 12.5 |
|  | 2H+50 μM DTHIB | 6.25 | 12.5 |
|  | FLC | 15.6 | NA |
|  | FLC+50 μM DTHIB | 15.6 | NA |
| *C. albicans* | 2H | 12.5 | 12.5 |
|  | 2H+25 μM DTHIB | 6.25 | 6.25 |
|  | 2H+50 μM DTHIB | 6.25 | 6.25 |
|  | FLC | 2 | NA |
|  | FLC+50 μM DTHIB | 2 | NA |

Notably, the reduction in MIC_90_ achieved by combining with DTHIB substantially improves the pharmacological profile of 2H by increasing the duration during which plasma concentrations are predicted to remain above the effective antifungal threshold based on our previously published pharmacokinetic analysis(67) (**Fig. S4**). Taken together, these findings indicate that interference with HSF1-dependent stress responses enhances the antifungal efficacy of Hsp110 inhibition.

### DTHIB similarly enhances the antifungal activity of 2H against Candida albicans but has little impact on fluconazole

To determine whether the interaction between DTHIB and 2H is conserved across fungal pathogens, we performed analogous studies using the widely used, clinically relevant *Candida albicans* strain SC5314. As observed in *C. neoformans*, DTHIB alone exhibited limited impact on growth, whereas co-treatment with DTHIB enhanced the antifungal activity of 2H (**Fig. 2C**). Analysis of antifungal susceptibility demonstrated that DTHIB reduced both the MIC_90_ and MFC of 2H, from 12.5 μM to 6.25 μM in the presence of DTHIB (**Table 1**). Although the magnitude of enhancement was somewhat less pronounced than that observed in *C. neoformans*, the overall pattern was similar.

Statistical analysis confirmed that DTHIB significantly potentiated the activity of 2H in *C. albicans*. Two-way ANOVA revealed significant effects of 2H concentration (F(7,72)=247.4, P<0.0001), DTHIB treatment (F(2,72)=14.69, P<0.0001), and a significant interaction between the two factors (F(14,72)=9.355, P<0.0001; **Table S2**). Bliss independence analysis further demonstrated synergistic interactions between 2H and DTHIB. Following Benjamini-Hochberg correction for multiple testing, significant positive Bliss scores were observed at 3.1 and 6.3 μM 2H in the presence of 50 μM DTHIB (**Fig. 2D; Table S5**). Similar but weaker synergistic interactions were observed with 25 μM DTHIB, where significant positive Bliss scores were detected only at 3.125 and 6.25 μM 2H (**Fig. S3B; Table S6**). Together, these findings indicate that inhibition of HSF1 enhances susceptibility to the Hsp110 inhibitor 2H in *C. albicans*, although the magnitude and concentration range of synergy are less pronounced than those observed in *C. neoformans*.

Fluconazole is the most commonly prescribed antifungal (89, 90). We therefore examined whether DTHIB enhances the antifungal activity of fluconazole. In contrast to its potentiating effect on 2H, DTHIB did not produce a biologically meaningful enhancement of fluconazole activity against either *C. neoformans* or *C. albicans* (**Fig. 3A,C**). Although two-way ANOVA detected statistically significant effects of treatment and treatment-concentration interactions, co-treatment with DTHIB did not alter the MIC_90_ of fluconazole in either species (**Table 1**), indicating that DTHIB does not enhance fluconazole susceptibility under these conditions.

**Figure 3.**
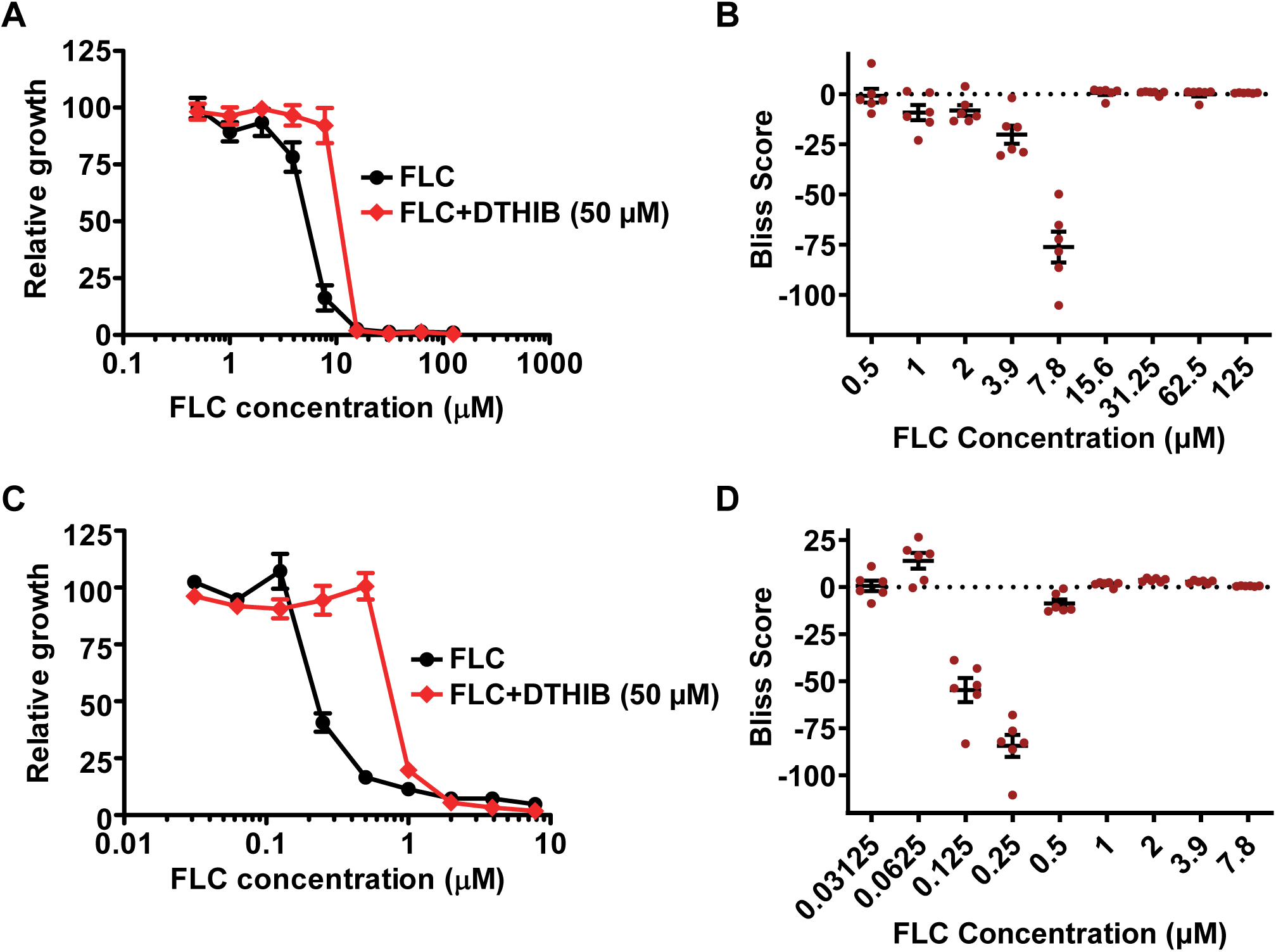
DTHIB does not enhance the antifungal activity of fluconazole. **(A,C)** Effect of DTHIB on the antifungal activity of fluconazole (FLC) against *C. neoformans* **(A)** and *C. albicans* **(C)**. Relative growth was normalized to the vehicle-treated control (100%). Data are presented as mean ± SEM from six independent biological replicates (n=6). Statistical significance was assessed by ordinary two-way ANOVA. **(B,D)** Bliss analysis of the data shown in **(A)** and **(C)**, respectively. Bliss scores were calculated using the Bliss independence model. Positive, zero, and negative Bliss scores indicate synergistic, additive, and antagonistic interactions, respectively. The dashed line indicates Bliss = 0. Data are presented as mean ± SEM (n = 6 for *C. neoformans* and n = 6 for *C. albicans*). Bliss scores were compared with the theoretical value of zero using a one-sample test, and P values were adjusted using the Benjamini-Hochberg false discovery rate (FDR) procedure. Detailed statistical results are provided in **Tables S7** and **S8**.

Consistent with these findings, Bliss analysis did not support a synergistic interaction between DTHIB and fluconazole (**Fig. 3B,D**). In *C. neoformans*, significant negative Bliss scores were observed at 3.9 and 7.8 μM fluconazole following Benjamini-Hochberg correction, indicating weak antagonistic interactions (**Table S7**). Although a statistically significant positive Bliss score was detected at 125 μM fluconazole, the magnitude of the effect was minimal (Bliss score = 0.58) and unlikely to represent biologically meaningful synergy. Similarly, in *C. albicans*, Bliss analysis revealed predominantly antagonistic interactions, with the strongest antagonism observed at 0.125 and 0.25 μM fluconazole (**Table S8**). While several higher concentrations yielded statistically significant positive Bliss scores after FDR correction, the magnitude of these effects was small (Bliss scores < 4) and not indicative of meaningful synergy. Together, these results indicate that DTHIB does not meaningfully enhance fluconazole activity and that its potentiating effect is relatively specific to Hsp110 inhibition by 2H. This specificity is consistent with the shared roles of HSF1 and Hsp110 in maintaining proteostasis and supporting fungal growth and survival.

### HSF1 inhibition accelerates the fungicidal activity of 2H

To further examine the functional consequences of simultaneously targeting HSF1 and Hsp110, we performed time-kill analyses using *C. neoformans* H99. Consistent with the lack of obvious impact on growth, DTHIB alone had little effect on fungal viability (**Fig. 4**). As shown in our previously published studies, 2H alone exhibited substantial fungicidal activity. However, fungal killing by 2H at 25 μM, the MFC, occurred gradually and required approximately 24 h to reduce viable cell numbers below the 99.9% killing threshold.

**Figure 4.**
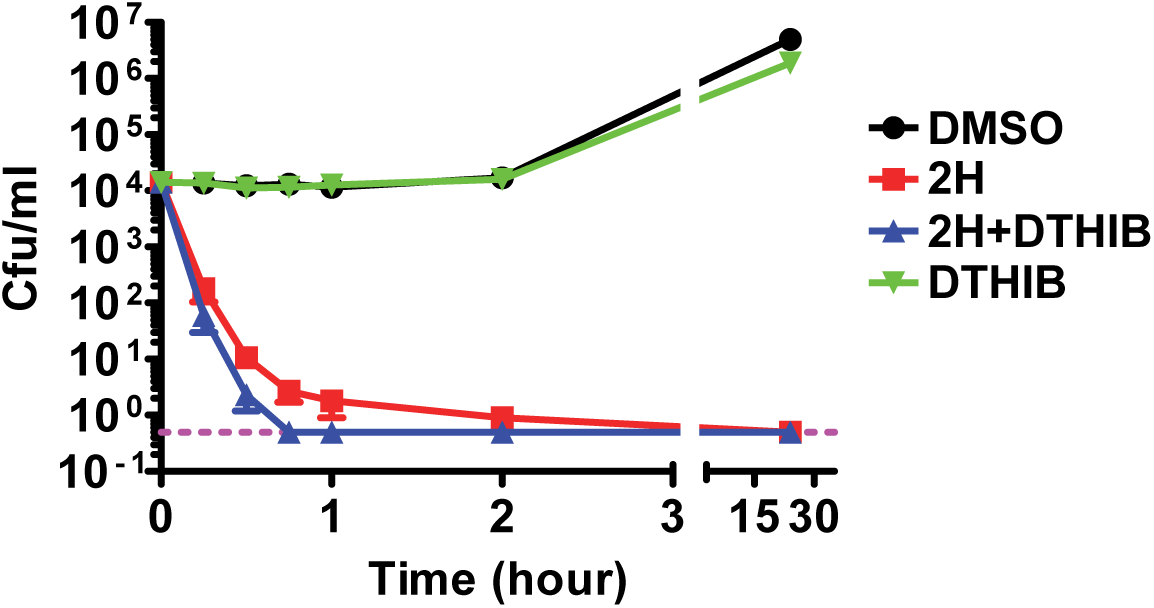
DTHIB enhances the fungicidal activity of 2H against *C. neoformans* while showing limited effect alone. Time-kill analysis of *C. neoformans* strain H99 treated with vehicle control, 2H (25 μM), DTHIB (50 μM), or the combination of 2H and DTHIB (2H+DTHIB). Viable cells were quantified by colony-forming unit (CFU) determination at the indicated time points. The dashed line indicates the limit of detection (>99.9% reduction relative to the starting inoculum). Data are presented as mean ± SEM from independent biological replicates (n=5).

In contrast, co-treatment with DTHIB dramatically accelerated fungal killing. The combination of 25 μM 2H and 50 μM DTHIB reduced viable cell numbers below the limit of detection (i.e., < 0.1% of the cells alive) within approximately 45 min. Thus, inhibition of HSF1 not only lowered the concentration of 2H required for fungicidal activity but also markedly increased the rate of fungal killing. These observations suggest that HSF1-dependent stress responses provide an important survival mechanism during Hsp110 inhibition.

### Inhibition of HSF1 enhances protein aggregation induced by Hsp110 inhibition in C. neoformans

Our published studies using *S. cerevisiae* and *C. albicans* demonstrated that treatment with 2H results in extensive accumulation of detergent-insoluble protein aggregates (66), consistent with the role of Hsp110 as a key regulator of proteostasis(28–44). We next investigated whether the enhanced antifungal activity of the combination treatment of 2H and DTHIB in *C. neoformans* was associated with increased protein aggregation. Detergent-insoluble protein aggregates were isolated from *C. neoformans* cells treated with vehicle, DTHIB, 2H, or the combination of 2H and DTHIB and subsequently analyzed by SDS-PAGE (**Fig. 5A**).

**Figure 5.**
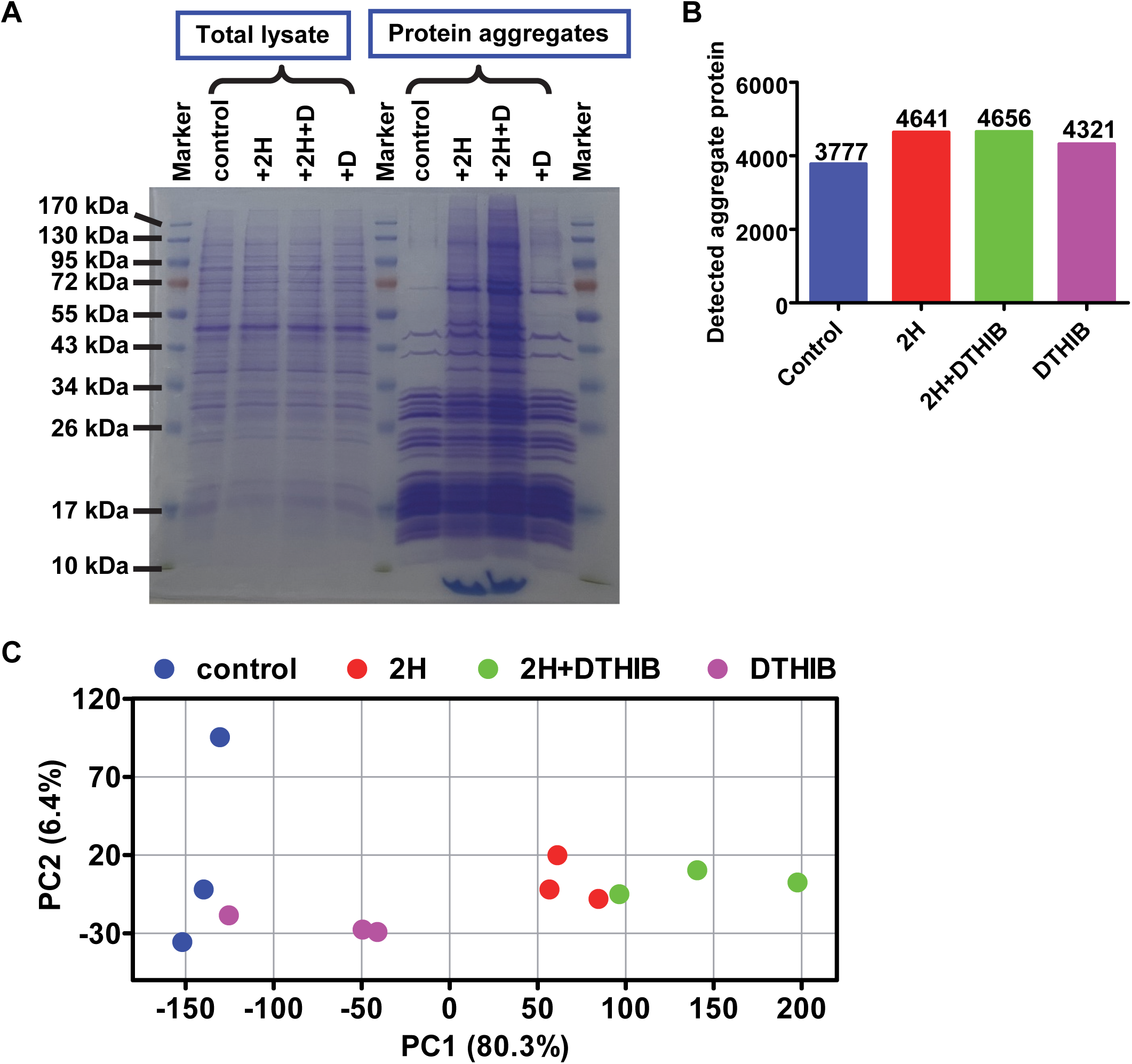
Hsp110 inhibition by 2H induces extensive protein aggregation in *C. neoformans,* and DTHIB further enhances this effect. **(A)** Analysis of protein aggregation *in vivo*. Detergent-insoluble protein aggregates were isolated from *C. neoformans* strain H99 treated with vehicle control, 2H (25 μM), DTHIB (D, 50 μM), or the combination of 2H and DTHIB (2H+D). Aggregated proteins were analyzed by SDS-PAGE. Total cellular lysates were used as controls (left). A representative gel is shown. **(B)** Number of Proteins identified in detergent-insoluble aggregates. Label-free quantitative mass spectrometry was performed on detergent-insoluble protein aggregates isolated from *C. neoformans* H99 treated with vehicle control, DTHIB, 2H, or the combination of 2H and DTHIB (2H+DTHIB), as described in (**A**). Three independent biological replicates were analyzed for each condition. **(C)** Principal component analysis (PCA) of aggregated proteins identified by quantitative mass spectrometry. Each point represents an independent biological replicate.

Consistent with the established role of Hsp110 in maintaining protein solubility, treatment with 2H caused extensive accumulation of detergent-insoluble proteins while having relatively little effect on total protein abundance (**Fig. 5A**). DTHIB alone produced only modest changes in aggregate levels, consistent with its limited impact on growth. In contrast, combined treatment with 2H and DTHIB further increased the abundance of aggregated proteins relative to treatment with 2H alone. Multiple protein bands were selectively enriched within the aggregate fractions following co-treatment, indicating enhanced accumulation of aggregation-prone proteins. These findings suggest that HSF1-dependent stress responses buffer the proteotoxic consequences of Hsp110 inhibition and that inhibition of HSF1 exacerbates the proteostasis defects induced by 2H-mediated Hsp110 inhibition.

### Quantitative proteomics reveals extensive proteome-wide aggregation following Hsp110 inhibition, and HSF1 inhibition broadens the aggregated proteome

To gain insight into the proteins that depend on Hsp110-mediated proteostasis, we performed label-free quantitative mass spectrometry on detergent-insoluble aggregate fractions isolated from *C. neoformans* cells treated with vehicle, DTHIB, 2H, or 2H plus DTHIB, as described in **Fig. 5A**. Proteomic analysis identified thousands of proteins within the aggregate fractions (**Fig. 5B, Table S9**). A total of 3,777 proteins were detected in the control samples, compared with 4,641 proteins following treatment with 2H and 4,656 proteins following co-treatment with 2H and DTHIB. In comparison, 4,321 proteins were identified in samples treated DTHIB alone. Thus, consistent with the SDS-PAGE results in **Fig. 5A**, these findings demonstrate that inhibition of Hsp110 substantially expands the aggregated proteome and that inhibition of HSF1 further enhances this effect.

Principal component analysis revealed clear separation among treatment groups, indicating substantial differences in aggregate composition following perturbation of proteostasis pathways (**Fig. 5C**). Samples treated with 2H clustered distinctly from control and DTHIB-treated samples, while the combination of 2H and DTHIB produced an additional shift in aggregate composition, consistent with enhanced proteome disruption.

To characterize the consequences of Hsp110 inhibition on the fungal proteome, we compared aggregate fractions isolated from cells treated with 2H with those from control cells. We carried out statistical analysis to determine the distribution of proteins exhibiting increased or decreased aggregation levels. The volcano plot revealed overwhelmingly widespread accumulation of proteins within detergent-insoluble aggregates following treatment with 2H (**Fig. 6A**), demonstrating that inhibition of Hsp110 results in extensive proteome instability. Of the 4,641 proteins detected in 2H-treated samples, 3,262 showed increased aggregation (Up; **Table 2**). In sharp contrast, no protein exhibited reduced aggregation (Down). Thus, this widespread aggregation observed in 2H-treated samples is consistent with the SDS-PAGE results shown in **Fig. 5A** and supports the essential role of Hsp110 in maintaining proteostasis.

**Figure 6.**
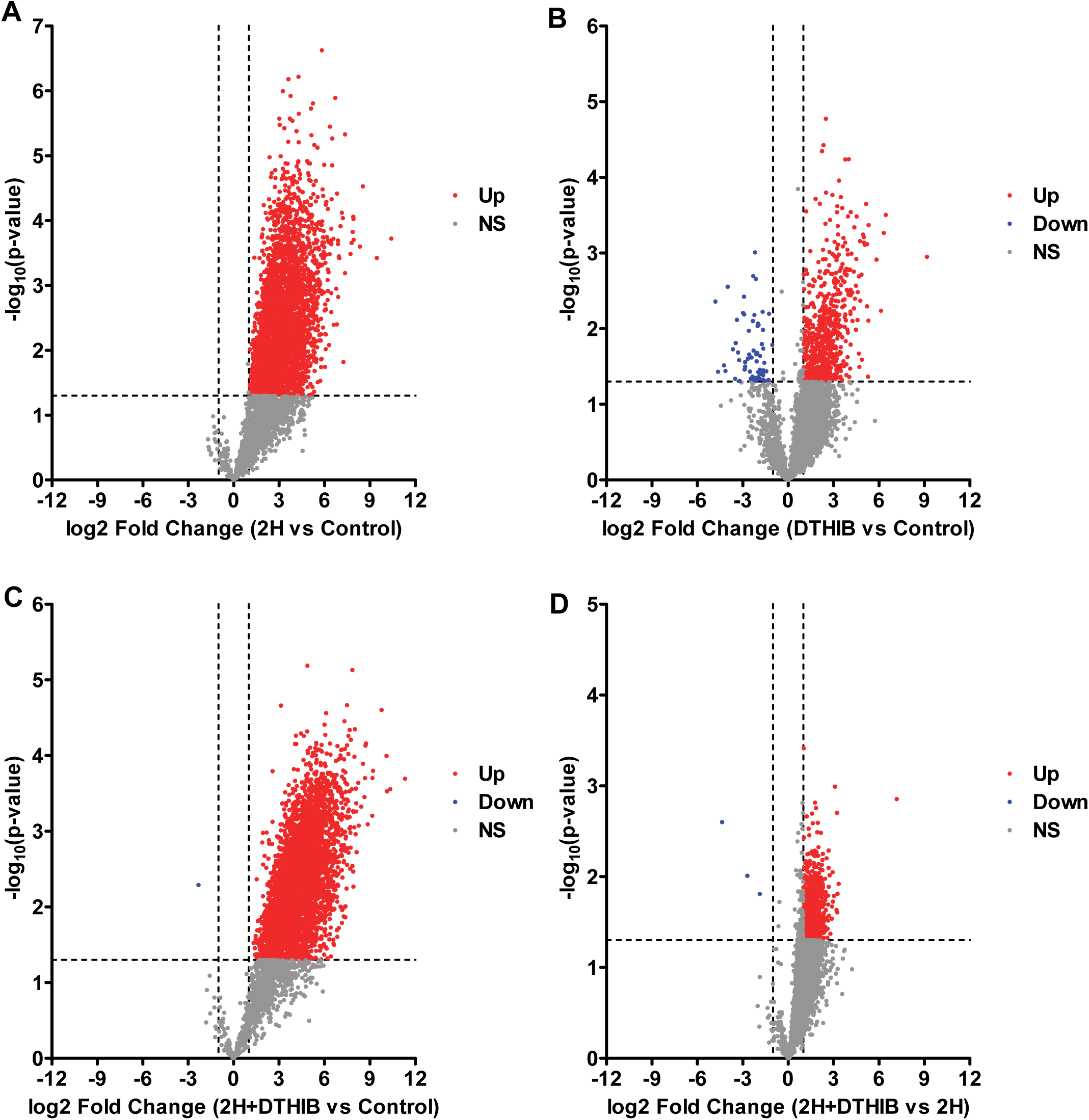
Aggregated proteome induced by 2H and DTHIB treatments. **(A-D)** The overall changes of detergent-insoluble proteins in *C. neoformans*. The volcano plots display the log₂ Fold Change in aggregate abundance as a function of statistical significance (−log_10_ *P*-value). Three independent biological replicates were obtained for each condition. Each point represents a protein identified in the aggregated fraction. Proteins exhibiting significantly increased and reduced abundance are highlighted in red and blue (Up, log₂ Fold Change ≥ 1 and *P* < 0.05; Down, log₂ Fold Change ≤ -1 and *P* < 0.05), respectively, whereas proteins not meeting these criteria are shown in gray (NS, not significantly changed). Dashed vertical lines indicate the log₂ Fold Change threshold at 1 and -1, and the dashed horizontal line indicates the significance threshold (*P* = 0.05). **(A)** 2H versus control, **(B)** combination of 2H and DTHIB (2H+DTHIB) versus control; **(C)** DTHIB versus control; **(D)** 2H+DTHIB versus 2H. Additional comparison of 2H+DTHIB versus DTHIB is shown in **Fig. S5**.

**Table 2.**
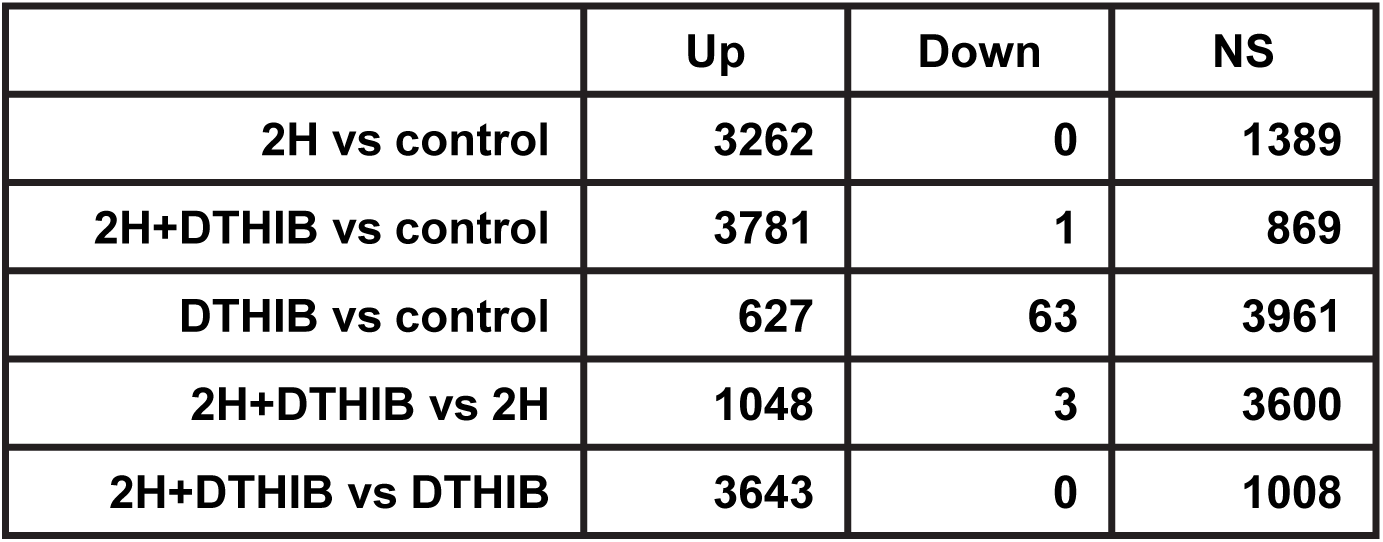
Comparison of the numbers of aggregated proteins identified among the treatments. The numbers of aggregated proteins in each category are obtained from the volcano plots in **Fig. 6** and **S5**. Up: log₂ Fold Change ≥ 1 and P-value < 0.05; Down: log₂ Fold Change ≤ -1 and P-value < 0.05; NS: not significantly changed. 2H+DTHIB: combined treatment of 2H and DTHIB.

|  | Up | Down | NS |
| --- | --- | --- | --- |
| 2H vs control | 3262 | 0 | 1389 |
| 2H+DTHIB vs control | 3781 | 1 | 869 |
| DTHIB vs control | 627 | 63 | 3961 |
| 2H+DTHIB vs 2H | 1048 | 3 | 3600 |
| 2H+DTHIB vs DTHIB | 3643 | 0 | 1008 |

In contrast to the extensive aggregation induced by 2H, DTHIB produced relatively limited changes in aggregate composition (**Fig. 6B** and **Table 2**), indicating that DTHIB alone is insufficient to induce widespread proteostasis disruption under these conditions. This observation is consistent with its minimal effects on cell growth and viability (**Fig. 2A** and **4**), as well as the limited aggregation observed by SDS-PAGE analysis (**Fig. 5A**). Of the 4,321 proteins detected in DTHIB-treated samples, 627 proteins showed enhanced aggregation (Up), while 63 proteins showed reduced aggregation (Down; **Table 2**). Although a measurable increase in protein aggregation was observed, the lack of detectable effects on growth or viability suggests that cells can tolerate these relatively modest disruptions in proteostasis. Interestingly, 617 of the 627 proteins showing increased aggregation also overlapped with proteins exhibiting increased aggregation in the 2H-treated samples (**Fig. S6A-B**), suggesting that these proteins may be particularly sensitive to disruption of chaperone function.

Notably, co-treatment of 2H and DTHIB induced additional aggregation relative to treatment with 2H alone (**Fig. 6C-D** and **Table 2**). Relative to the control, 3,781 proteins showed enhanced aggregation (Up), whereas only a single protein exhibited reduced aggregation (Down). Moreover, compared with the 2H-treated samples, 1,048 proteins showed increased aggregation among the 4,656 proteins detected in the co-treatment samples (**Table 2**). Among proteins exhibiting increased aggregation relative to the control, the 2H-treated and co-treatment groups share 3,186 overlapping proteins, representing 97.7% of the proteins exhibiting increased aggregation in the 2H-treated samples (**Fig. S6A-B**). Thus, most proteins enriched in aggregate fractions during co-treatment overlapped with proteins aggregated following 2H treatment alone, indicating that Hsp110 inhibition generates a core aggregation program. However, additional proteins accumulated in aggregates following the addition of DTHIB, resulting in a further expansion of the aggregated proteome.

Together, these findings indicate that HSF1-dependent stress responses help buffer the proteotoxic consequences of Hsp110 inhibition. Inhibition of HSF1 broadens the range of proteins that lose solubility during Hsp110 inhibition, resulting in enhanced proteome instability and providing a mechanistic explanation for the potentiated antifungal activity observed during co-treatment with DTHIB and 2H.

### Hsp110 inhibition induces widespread aggregation of high-molecular-weight proteins across diverse functional protein classes and cellular pathways

To dissect the common features of the proteins affected by Hsp110 inhibition following 2H treatment, we next examined whether protein size influences susceptibility to aggregation. First, the mean molecular weight and the average molecular weight within each quartile were calculated for proteins showing enhanced aggregation (Up) (**Table 3**). For comparison, proteins with no significant changes (NS) were used. The mean molecular weights for the Up group were significantly higher than those of the NS group. Combined treatment with DTHIB further increased the molecular weight of proteins exhibiting elevated aggregation levels (**Table S10**).

**Table 3.** Comparison of molecular weight in aggregated proteins following 2H treatment. Molecular weights for mean, median, 25^th^ and 75^th^ percentile are shown for the NS and Up populations of the aggregated proteins induced by 2H treatment versus the control identified in the volcano plot in **Fig. 6A**. Up: log₂ Fold Change ≥ 1 and P-value < 0.05; NS: not significantly changed.

| 2H vs control | Mean (Da) | Median (50<sup>th</sup> percentile) (Da) | 25<sup>th</sup> percentile (Da) | 75<sup>th</sup> percentile (Da) |
| --- | --- | --- | --- | --- |
| NS | 55,000 | 43,333 | 27,829 | 73,248 |
| Up | 63,234 | 53,070 | 35,502 | 79,126 |

Next, we divided proteins with increased aggregation into six equal groups according to molecular weight, and aggregate enrichment was calculated as log_2_ Fold Change (log_2_ FC; **Table 4**). The lower half of the molecular weight distribution (S1-S3, <53 kDa) is relatively flat, with median log_2_ FC values ranging from 2.93 to 2.96. Starting at approximately 53 kDa (S4), the median log_2_ FC increased steadily. The highest molecular weight group (>93 kDa) exhibited the highest median log_2_ FC value (3.59), about 22% greater than that of the lowest molecular weight group (3.59 vs 2.96). These results suggest that the effect of molecular weight on aggregation is driven primarily by proteins larger than approximately 50-70 kDa, whereas proteins below ∼50 kDa display similar aggregation behavior.

**Table 4.** Hsp110 inhibition by 2H induces widespread protein aggregation with high molecular weight. Protein molecular weight is associated with extend of aggregation following Hsp110 inhibition by 2H. The Up population (log₂ Fold Change ≥ 1 and P-value < 0.05) identified in the volcano plot for 2H vs control in **Fig. 6A** were divided into six groups based on molecular weight with roughly equal number of proteins in each group. Aggregate enrichment was calculated as median log₂ Fold Change (FC).

| Group | Molecular Weight Range (kDa) | Numbers of Proteins | Median log2 FC |
| --- | --- | --- | --- |
| S1 | < 29.10 | 529 | 2.958 |
| S2 | 29.11 - 40.52 | 528 | 2.933 |
| S3 | 40.53 - 53.07 | 528 | 2.945 |
| S4 | 53.08 - 68.69 | 528 | 3.109 |
| S5 | 68.72 - 93.30 | 528 | 3.321 |
| S6 | > 93.46 | 529 | 3.591 |

Taken together, these findings indicate that larger and structurally more complex proteins are particularly vulnerable to loss of Hsp110 function. This observation is consistent with previous studies in *S. cerevisae* showing that deletion of the major Hsp110 homolog *SSE1* results in the aggregation of high-molecular-weight proteins (91). These findings suggest that Hsp110 most likely plays an important role in assisting the folding of large, multi-domain proteins.

Functional classification revealed that proteins enriched within aggregate fractions following Hsp110 inhibition participate in a broad range of essential cellular processes (**Fig. 7A** for GO biological processes and **Fig. S7** for KEGG pathways). Prominent categories included various metabolic processes (such as protein, nucleic acid, and small molecules metabolism), cellular localization, gene expression, intracellular transport, the spliceosome, and the proteasome. In addition, these proteins are distributed across diverse cellular compartments (**Fig. 7B**) and possess a wide range of molecular functions (**Fig. S8**). Consistent with the established role of Hsp110 as a major molecular chaperone in the cytosol and nucleus, proteins localized to these compartments constituted the majority of those enriched within aggregate fractions following 2H treatment. The enrichment of proteins associated with intracellular organelles and protein-containing complexes further supports a role for Hsp110 in facilitating protein transport across various membranes and promoting protein complex assembly. Interestingly, mitochondrial proteins were among the most highly enriched categories, consistent with the proposed function of Hsp110 in facilitating protein import into mitochondria. Similar patterns were observed following co-treatment of 2H and DTHIB.

**Figure 7.**
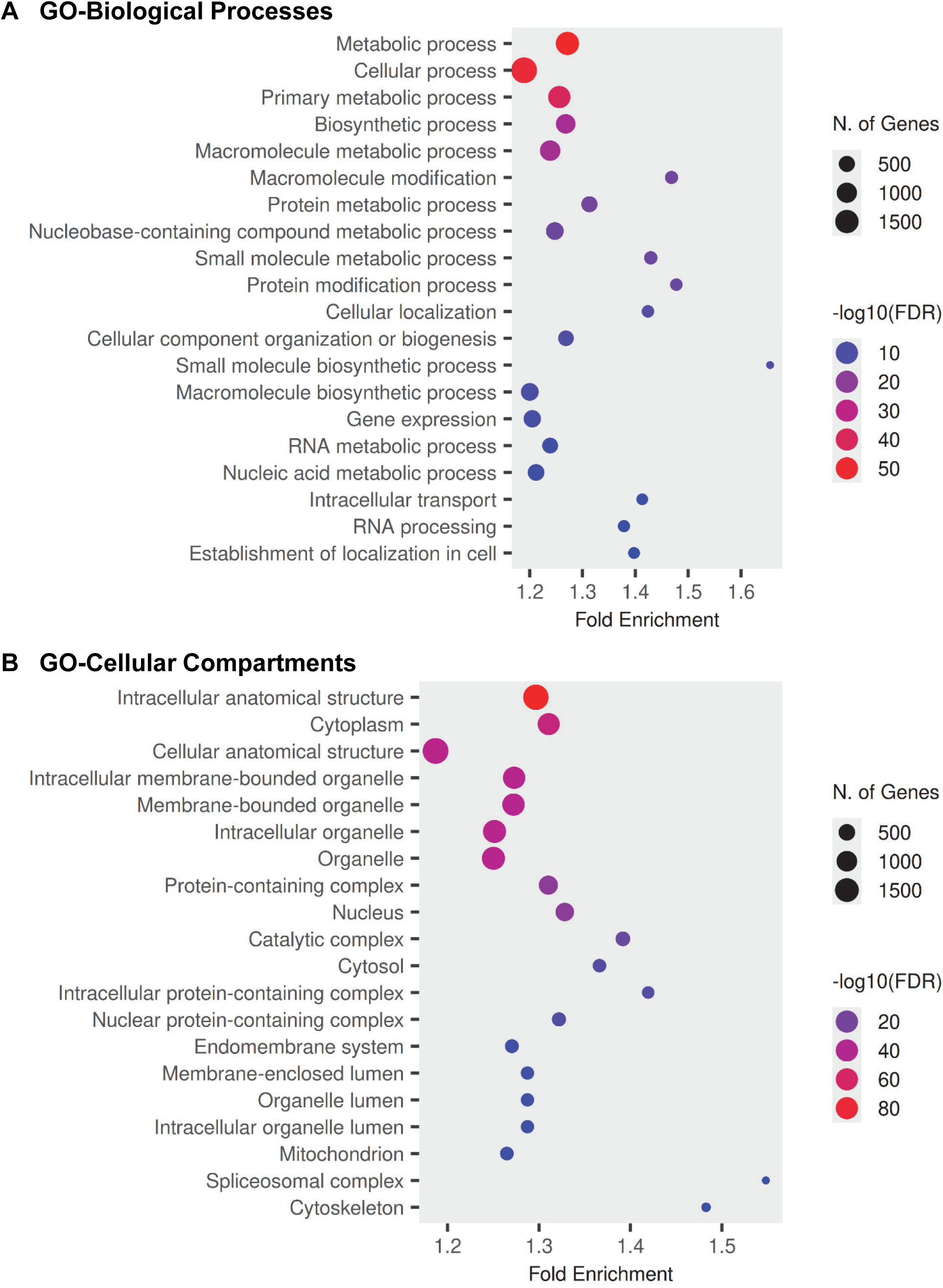
The top 20 GO biological processes (A) and cellular compartments (B) enriched in the aggregated proteome of *C. neoformans* induced by 2H treatment. The plots were generated using ShinyGO 8.6 (South Dakota State University).

Notably, more than a dozen virulence-associated proteins accumulated in the aggregate fractions of samples treated with 2H (**Table S11**), including components of the calcineurin signaling pathway (Cbp1 and cyclophilin)(92, 93). Furthermore, the aggregation of most of these proteins increased further upon co-treatment with DTHIB. These findings suggest that 2H, particularly when combined with DTHIB, has the potential to attenuate virulence in *C. neoformans*. Taken together, these observations demonstrate that Hsp110 contributes broadly to proteome maintenance and that its inhibition causes widespread disruption of cellular homeostasis.

## DISCUSSION

The increasing prevalence of invasive fungal infections and the emergence of antifungal resistance highlight the need for therapies that target biological processes distinct from those exploited by current antifungal agents. In this study, we demonstrate that inhibition of HSF1 significantly potentiates the antifungal activity of the fungal Hsp110 inhibitor 2H. Co-treatment with DTHIB and 2H reduced both the MIC_90_ and MFC of 2H, accelerated fungal killing, and enhanced protein aggregation in *C. neoformans*. Proteomic analyses further revealed that inhibition of HSF1 amplifies the proteostasis defects caused by Hsp110 inhibition, leading to widespread disruption of protein solubility across the fungal proteome. Together, these findings establish a functional connection between HSF1-mediated stress responses and Hsp110-dependent proteostasis and support simultaneous targeting of these pathways as a promising novel antifungal strategy.

Hsp110 represents an attractive antifungal target because of its central role in maintaining protein homeostasis. Unlike many existing antifungal agents that inhibit specific biosynthetic pathways, primarily the cell membrane and cell wall (9, 13–15), Hsp110 regulates protein folding, prevents aggregation, and maintains proteome integrity (28–44). Consistent with these functions, inhibition of Hsp110 by 2H induced widespread aggregation of proteins involved in diverse cellular processes. These findings indicate that Hsp110 acts as a global regulator of proteome stability in fungal cells and that its inhibition simultaneously compromises multiple essential cellular pathways, providing a mechanistic explanation for the potent antifungal activity of 2H.

An important implication of this study extends beyond antifungal drug discovery and addresses a long-standing question in Hsp110 biology. Despite extensive investigation, the endogenous proteins that depend on Hsp110 chaperone activity remain poorly defined because Hsp110–protein interactions are believed to be highly dynamic and transient. The unique mechanism of action of 2H provides a powerful approach for addressing this challenge. Proteins that lose solubility and accumulate in detergent-insoluble aggregates following Hsp110 inhibition are likely to depend, directly or indirectly, on Hsp110-mediated proteostasis. Thus, the aggregate proteome generated following 2H treatment provides a valuable starting point for future studies aimed at identifying Hsp110-dependent proteins and understanding the biological functions of Hsp110 *in vivo*.

Several observations from our proteomic analyses provide clues regarding the characteristics of proteins that rely on Hsp110 activity. Aggregate enrichment increased progressively with protein molecular weight, suggesting that large and structurally complex proteins are particularly vulnerable to loss of Hsp110 function. Although the current data do not distinguish direct Hsp110 substrates from proteins that aggregate secondarily following proteostasis collapse, they provide an important foundation for future studies aimed at defining endogenous Hsp110-dependent proteins and elucidating how Hsp110 maintains proteome integrity. Another interesting feature is the enrichment of organelle proteins in the aggregation. This is consistent with the proposed role of Hsp110 in protein transport across the membranes such as protein import into ER and mitochondria. While the role of Hsp110 in protein import into the ER has been widely recognized and experimentally supported, there is little evidence to support the involvement of Hsp110 in protein import into mitochondria despite being proposed. The enrichment of mitochondrial proteins (**Fig. 7B**) provides support for Hsp110’s role in protein import into mitochondria.

Our results also reveal an important protective role for HSF1-mediated stress responses during proteostasis disruption. HSF1 coordinates the expression of molecular chaperones and protein quality-control factors that enable cells to adapt to proteotoxic stress (68–79). Thus, HSF1 occupies a central position in the proteostasis network. Although DTHIB alone produced only modest effects on fungal growth and protein aggregation, its impact became pronounced when Hsp110 function was compromised. These observations suggest that HSF1 functions as a buffering mechanism that helps fungal cells withstand the proteotoxic stress generated by Hsp110 inhibition. Simultaneous disruption of both pathways therefore overwhelms the cellular capacity to maintain protein homeostasis, resulting in extensive proteome instability and rapid loss of viability.

Importantly, our findings provide experimental support for a long-standing hypothesis in fungal biology. HSF1 is essential for growth, stress adaptation, and virulence in multiple fungal species and has therefore been proposed as a promising target for antifungal drug development (80–82). However, direct evidence supporting pharmacological inhibition of HSF1 as an antifungal strategy has remained limited, largely because of the lack of suitable approaches for inhibiting fungal HSF1. To our knowledge, this study provides the first pharmacological evidence that inhibition of HSF1-dependent stress responses can effectively sensitize fungal pathogens to proteotoxic stress. The positive Bliss scores observed across multiple concentrations further support a cooperative interaction between HSF1 and Hsp110 inhibition rather than a merely additive effect. The ability of DTHIB to enhance the antifungal efficacy of 2H specifically, accelerate fungal killing, and exacerbate proteostasis collapse supports HSF1 as a promising antifungal sensitization target and establishes proof-of-concept for therapeutically targeting fungal HSF1.

There are at least two possible explanations for the relatively modest activity of DTHIB as a single agent. First, DTHIB was originally discovered and optimized as an inhibitor of human HSF1 rather than fungal HSF1. Although fungal HSF1 proteins exhibit sequence conservation relative to their human counterpart, DTHIB may bind fungal HSF1 with reduced affinity, resulting in incomplete target inhibition. Second, fungal cells possess unique structural barriers that can limit intracellular drug accumulation. The thick polysaccharide cell wall and ergosterol-containing plasma membrane may reduce uptake and retention of DTHIB, thereby decreasing effective intracellular target engagement. These factors may explain why DTHIB alone produces only limited effects despite the essential role of HSF1 in fungal biology. Nevertheless, the pronounced enhancement of 2H-mediated antifungal activity strongly argues that HSF1-dependent stress responses constitute an important protective mechanism during proteostasis disruption.

The present study provides a clear direction for future therapeutic development. An important next step will be the development of inhibitors with improved specificity and potency toward fungal HSF1. Structure-guided medicinal chemistry aimed at increasing affinity for fungal HSF1 while improving penetration through the fungal cell wall and membrane could substantially enhance antifungal efficacy and further strengthen the synergy observed with Hsp110 inhibition. Such compounds would not only represent improved antifungal leads but would also provide powerful tools for dissecting fungal stress-response networks and HSF1 biology.

In summary, our study identifies HSF1-mediated stress responses as a critical defense mechanism against proteotoxic stress induced by Hsp110 inhibition. Fungal cells rely on HSF1-dependent stress responses to withstand the proteome damage caused by loss of Hsp110 function, and simultaneous disruption of both pathways triggers extensive proteome instability and enhanced antifungal activity. Beyond establishing dual targeting of HSF1 and Hsp110 as a promising antifungal strategy, our work demonstrates the utility of 2H as a chemical probe for studying Hsp110-dependent proteostasis and provides a foundation for future investigations into the endogenous proteins and biological processes that rely on Hsp110 activity. These findings advance both antifungal drug discovery and our understanding of Hsp110 biology.

## MATERIALS AND METHODS

### *In silico* docking of DTHIB to HSF1 proteins

The DNA-binding domains (DBDs) of human HSF1 (hHSF1) and *C. neoformans* HSF (cnHSF1) were compared to evaluate DTHIB binding. The DNA-bound hHSF1 structure (PDB code: 5D5V) served as the reference. The cnHSF1 DBD (residues 78-186) was modeled from its amino acid sequence using AlphaFold. Sequence alignment with Clustal 2.1 showed 51.9% identity and conservation of key HSF1 DBD motifs. The modeled cnHSF1 DBD was structurally aligned to the two DNA-bound hHSF1 DBDs in 5D5V, generating a DNA-bound cnHSF1 model while retaining the DNA duplex for subsequent analyses.

Both the hHSF1 and cnHSF protein-DNA complexes were prepared using the Schrödinger Protein Preparation workflow. Hydrogen atoms were added, bond orders assigned, and protonation states optimized. Potential DTHIB-binding pockets were identified with SiteMap, which evaluates cavity geometry and physicochemical properties (94). DNA was retained during SiteMap analysis to identify pockets accessible in the DNA-bound state. Pockets outside the protein-DNA interface were selected for docking.

DTHIB was prepared by adding polar hydrogens, assigning Gasteiger charges, defining rotatable bonds, and generating a PDBQT file. Docking was performed with AutoDock Vina v1.2.7 (95, 96). SiteMap pocket centroids defined the search space, with the receptor treated as rigid and DTHIB as flexible. Docking parameters were identical for hHSF1 and cnHSF1 (exhaustiveness = 64, maximum modes = 20, energy range = 5 kcal mol⁻¹) to enable direct comparison. Docking poses were ranked by the Vina scoring function and inspected in PyMOL, and poses that substantially overlapped the DNA-binding interface were excluded.

Protein-ligand interactions were analyzed using PLIP, which identifies hydrogen bonds, hydrophobic contacts, π-mediated interactions, salt bridges, and other noncovalent interactions (97, 98). Interaction networks were visualized in PyMOL using identical representations for hHSF1 and cnHSF1 to facilitate comparison. Conservation of the DTHIB-binding environment was assessed using both sequence and structural information rather than residue numbering alone.

### Antifungal susceptibility and viability testing on *C. neoformans* Type Strain H99 and *Candida albicans* strain SC5314

Overall, MIC and MFC determinations were performed according to the Clinical and Laboratory Standards Institute for yeasts (CLSI, M27) and our previously published protocol (66), with slight modifications.

For *C. neoformans*, the Type Strain H99 was obtained from the laboratory of Dr. Joseph Heitman (Duke University). After growing in Sabouraud medium, this strain was resuspended in 35% glycerol, aliquoted, flash frozen in liquid nitrogen, and stored in a −80 °C freezer.

For antifungal susceptibility tests, a frozen aliquot of H99 was thawed and streaked on agar plates containing Sabouraud medium. The plates were incubated at 30 °C for 2-3 days to allow single colonies to grow. Single colonies were inoculated into a 5-ml Sabouraud liquid medium and grown at 30 °C for 1 overnight. The culture was pelleted by centrifugation at 1,200 g for 5 min at room temperature and washed twice using PBS at room temperature. The inoculum was prepared using the RPMI medium with 5×10^3^-1×10^4^ colony forming unit (CFU) (corresponding to an OD_600_ of 0.001). Sterile 96-well culture plates were used for growth test. After aliquoting 100 µl of the inoculum into each well, serial dilutions of 2H, fluconazole, or DTHIB alone or in combination were added to the wells and mixed well by pipetting up and down. After shaking at 37 °C for 48 hours, growth was measured by absorbance at 600 nm using a spectrophotometric plate reader. Wells with only medium were used as blanks and wells with only inoculum were used as growth controls. A viability test to determine the minimum fungicidal concentration (MFC) involved plating aliquots from wells with no observable growth on plates containing Sabouraud agar, followed by colony counting after incubation.

A similar protocol was used for testing *C. albicans*. The WT *C. albicans* strain SC5314 was obtained from Dr. Ronda Rolfes. RPMI medium was used (99). Briefly, after growing at 30 °C for 1 overnight, the culture was centrifuged at 1,200 g for 5 min at room temperature and washed twice with PBS. The inoculum was prepared by diluting the washed culture into RPMI medium to a final OD_600_ of 0.0005 or 5×10^3^-1×10^4^ colony forming unit (CFU) per ml. The inoculum was aliquoted into a sterile 96-well culture plate. Serial dilutions of compounds were prepared and added into the aliquoted inoculum. The plate was incubated at 37 °C for 24 hours. The growth was measured using MTT assay with absorbance at 510 nm in a spectrophotometric plate reader. Wells with medium only were used as blanks and wells with only inoculum were used as growth controls. As a viability test to determine MFC, a 200 µl aliquot was plated on to a Sabouraud agar plate. After growing at 30 °C for 1-2 days, colonies were counted.

Bliss synergy scores were calculated using the Bliss independence model. Expected relative growth was calculated as the product of the relative growth observed for each single agent alone. The Bliss score was defined as expected relative growth minus observed relative growth for the combination treatment. Relative growth values were first normalized to vehicle-treated controls prior to Bliss analysis. Positive scores indicate synergy, scores near zero indicate additivity, and negative scores indicate antagonism.

Statistical analyses were performed using GraphPad Prism. For growth inhibition assays, differences among treatment groups were analyzed using two-way ANOVA followed by Sidak’s multiple-comparison test. Bliss synergy scores were analyzed using a one-sample Wilcoxon signed-rank test against a theoretical value of zero (additivity), with correction for multiple comparisons where applicable. P values <0.05 were considered statistically significant.

### Time-kill analysis on *C. neoformans* Type Strain H99

A single colony of H99 was inoculated into a 5 ml Sabouraud medium. After growing at 30 °C for 1 overnight, the culture was pelleted and washed twice using PBS at room temperature. The inoculum was prepared using the RPMI medium with 5×10^3^-1×10^4^ colony forming unit (CFU) (corresponding to an OD_600_ of 0.001). A 5-ml culture was grown at 37 °C with addition of 2H, DTHIB, or the combination. DMSO, the solvent for 2H and DTHIB, was used as a negative control. At the indicated time points, a 200 µl aliquot was taken and plated on to a Sabouraud agar plate. After growing at 30 °C for 2-3 days, colonies were counted.

### *In vivo* protein aggregation assay

The published protocols for analyzing protein aggregation were used (91, 100). A single colony of the *C. neoformans* Type Strain H99 was inoculated into Sabouraud medium. After growing one overnight, the culture was centrifuged, washed with PBS. Then it was diluted to 0.2 OD_600_ in RPMI and continued to grow for another 3-4 hours to reach 0.4-0.5 OD_600_. 2H or DTHIB was added to a final concentration of 25 and 50 µM, respectively. Samples were taken after a 90- min treatment. Each sample was centrifuged at 1,200 g for 5 min to pellet the cells and flash frozen in liquid nitrogen. The cell pellets were thawed and resuspended in a cold PBS buffer with addition of 1 mM EDTA, 2 mM DTT and 0.1% Tween 20. French Press was applied to break the cells. Intact cells and cell debris were removed by a centrifugation at 5,000 g for 10 min. Protein concentrations of the cell lysates were estimated by OD_280_ using NanoDrop. After adjusting the concentrations to a similar level, the cell lysates were subjected to an ultracentrifugation at 20,000 g for 20 min. The supernatant was removed, and the pellets were washed with a cold PBS buffer containing 1 mM EDTA, 2 mM DTT, and 2% NP40. After a second centrifugation at 20,000 g for 20 min, the resulting pellets were dissolved in lithium dodecyl sulfate (LDS) sample buffer. The total cell lysate and pellet for each treatment were loaded onto an SDS-PAGE. Equivalent amount of each sample was adjusted based on the OD_280_ measurement.

### Mass spectrometry analysis

Sample preparation: After normalization to the corresponding total lysate, aggregated proteins were run on SDS-PAGE for ∼10 min to allow all the proteins enter the separating gel for 2-3 mm. Then, the electrophoresis was stopped, and SDS-PAGE was stained with Coomassie blue. After de-staining, the gel was soaked in ddH_2_O. The protein bands were excised from the gel, cut into equal size cubes approximately 1mm, and transferred to an Eppendorf tube. The gel pieces were washed with ddH_2_O for 10 min. Further processing was completed using the Preomics reagents as follows with slight modification. The gel pieces were incubated with 50 µl LYSE solution at 75 °C with continuous mixing at 1,000 rpm for 10 min in a Thermo mixer. The gel pieces were cooled and incubated with 25 µl DIGEST for 2 hours at 37 °C with continuous mixing at 1,000 rpm. STOP solution (100 µl) was added. The solution was transferred to the Preomics spin cartridge and spun for 2 min at 3,800 g. WASH 1 (200 µl) was added to the cartridge and spun at 3,800 g for 2 min. WASH 2 (200 µl) was added to the cartridge and spun at 3,800 g for 2 min. The peptides were eluted and collected in a fresh Eppendorf tube by adding 100 µl ELUTE to the cartridge and spinning at 3,800 g for 2 min. The elution step was repeated. The 200 µl total of digested peptides were collected into fresh tubes for subsequent MS analysis.

LC-MS/MS: LC-MS/MS analysis were performed using an Orbitrap Lumos Tribrid (Thermo) mass spectrometer coupled to an Easy nLC 1200 (Thermo) nanoflow UPLC system. The LC-MS/MS system was fitted with a Flex ion source (Thermo) and series RSLC 75µm x 50cm with an Aurora XT 75umx250mm C18 1.7um (Ion Optick). The mobile phase consists of Buffer A (0.1% formic acid in water) and Buffer B (80% acetonitrile in water and 0.1% formic acid). The peptides were injected onto the above column assembly and eluted with an acetonitrile/0.1% formic acid gradient at a flow rate of 300 nL/min over 1 hour. The nano-spray ion source was operated at 1.9 kV. The digests were analyzed using a data independent acquisition (DIA) method acquiring a full scan mass spectrum (350-1200m/z), at an AGC = 300% and an ITmax = Auto followed by targeted MS_2_ analysis with 46 Da window. NCE = 30%, AGC = Custom 1000%, and ITmax = Auto.

Data Analysis: The data were analyzed in Spectronaut (ver 20.1) using the Pulsar search algorithm and the database for *C. neoformans*. Proteins were identified at an FDR <0.01 and quantification used the MS2 level peptide intensities.

Perseus (version 2.1.3.0) (101) was used for standard volcano plot analyses, and the plots were generated using GraphPad Prism. Functional enrichment analysis was performed using ShinyGO 0.86 (South Dakota State University) (102). Significantly enriched Gene Ontology (GO) Biological Process (BP), Cellular Component (CC), Molecular Function (MF), and KEGG pathway terms were identified.

### Statistics and Reproducibility

All experiments were performed with at least three independent biological replicates. Exact n values are provided in figure legends. Data are presented as the mean ± standard deviation unless otherwise indicated.

### Data Availability

All data supporting the findings of this study are included in the manuscript and its supplementary information files. Source data are provided with this paper. The proteomics datasets generated during this study will be deposited in the PRIDE repository and made publicly available.

## ACKNOWLEDGEMENTS

We thank Dr. Joseph Heitman (Duke University) for the *C. neoformans* Type Strain H99 and Dr. Ronda Rolfes (Georgetown University) for the *C. albicans* strain SC5314. We thank Anusri Arun Prassath, Bao-Tich Nguyen, Spoorti Kalashetti, Nancy Zou, Aseel Packry, and Claudia Romero Carpio for their technical assistance and valuable contributions to this work. We further thank graduate student Ayil Perrott for critically reading and editing the manuscript.

This work was supported by the National Institute of General Medical Sciences of National Institutes of Health (R01GM098592 to Q.L.); Virginia Commonwealth University Clinical and Translational Science Award (UL1TR002649) from the National Center for Advancing Translational Sciences and the CCTR Endowment Fund of Virginia Commonwealth University (CCTR Award to Q.L.); the Commonwealth Health Research Board Funding (CHRB Award to Q.L.); and VCU Massey Comprehensive Cancer Center Proteomics Shared Resource, supported, in part, with funding from NIH-NCI Cancer Center Support Grant P30 CA016059. The content is solely the responsibility of the authors and does not necessarily represent the official views of the National Institutes of Health.

## Notes

### Competing Interest Statement

The authors have declared no competing interest.

## REFERENCES

1. High time to tackle drug-resistant fungal infections. Nature. 2025;640(8059):569.

2. Brown GD, Denning DW, Gow NA, Levitz SM, Netea MG, White TC. Hidden killers: human fungal infections. Sci Transl Med. 2012;4(165):165rv13.

3. Kullberg BJ, Arendrup MC. Invasive Candidiasis. N Engl J Med. 2015;373(15):1445–56.

4. Magill SS, O’Leary E, Janelle SJ, Thompson DL, Dumyati G, Nadle J, et al. Changes in Prevalence of Health Care-Associated Infections in U.S. Hospitals. N Engl J Med. 2018;379(18):1732–44.

5. Pfaller MA, Diekema DJ. Epidemiology of invasive candidiasis: a persistent public health problem. Clin Microbiol Rev. 2007;20(1):133–63.

6. Benedict K, Jackson BR, Chiller T, Beer KD. Estimation of Direct Healthcare Costs of Fungal Diseases in the United States. Clin Infect Dis. 2019;68(11):1791–7.

7. Schille TB, Sprague JL, Naglik JR, Brunke S, Hube B. Commensalism and pathogenesis of Candida albicans at the mucosal interface. Nat Rev Microbiol. 2025;23(8):525–40.

8. Lohse MB, Gulati M, Johnson AD, Nobile CJ. Development and regulation of single- and multi-species Candida albicans biofilms. Nat Rev Microbiol. 2018;16(1):19–31.

9. Costa-de-Oliveira S, Rodrigues AG. Candida albicans Antifungal Resistance and Tolerance in Bloodstream Infections: The Triad Yeast-Host-Antifungal. Microorganisms. 2020;8(2).

10. Idnurm A, Bahn YS, Nielsen K, Lin X, Fraser JA, Heitman J. Deciphering the model pathogenic fungus Cryptococcus neoformans. Nat Rev Microbiol. 2005;3(10):753–64.

11. Chang CC, Harrison TS, Bicanic TA, Chayakulkeeree M, Sorrell TC, Warris A, et al. Global guideline for the diagnosis and management of cryptococcosis: an initiative of the ECMM and ISHAM in cooperation with the ASM. Lancet Infect Dis. 2024;24(8):e495–e512.

12. Lin X, Heitman J. The biology of the Cryptococcus neoformans species complex. Annu Rev Microbiol. 2006;60:69–105.

13. Salazar SB, Simoes RS, Pedro NA, Pinheiro MJ, Carvalho M, Mira NP. An Overview on Conventional and Non-Conventional Therapeutic Approaches for the Treatment of Candidiasis and Underlying Resistance Mechanisms in Clinical Strains. J Fungi (Basel). 2020;6(1).

14. Berman J, Krysan DJ. Drug resistance and tolerance in fungi. Nat Rev Microbiol. 2020;18(6):319–31.

15. Murphy SE, Bicanic T. Drug Resistance and Novel Therapeutic Approaches in Invasive Candidiasis. Front Cell Infect Microbiol. 2021;11:759408.

16. Chen SC, Sorrell TC. Antifungal agents. Med J Aust. 2007;187(7):404–9.

17. Lee A. Ibrexafungerp: First Approval. Drugs. 2021;81(12):1445–50.

18. Carolus H, Pierson S, Lagrou K, Van Dijck P. Amphotericin B and Other Polyenes-Discovery, Clinical Use, Mode of Action and Drug Resistance. J Fungi (Basel). 2020;6(4).

19. Balch WE, Morimoto RI, Dillin A, Kelly JW. Adapting proteostasis for disease intervention. Science. 2008;319(5865):916–9.

20. Labbadia J, Morimoto RI. The biology of proteostasis in aging and disease. Annu Rev Biochem. 2015;84:435–64.

21. Balchin D, Hayer-Hartl M, Hartl FU. In vivo aspects of protein folding and quality control. Science. 2016;353(6294):aac4354.

22. Klaips CL, Jayaraj GG, Hartl FU. Pathways of cellular proteostasis in aging and disease. J Cell Biol. 2018;217(1):51–63.

23. Chen B, Retzlaff M, Roos T, Frydman J. Cellular strategies of protein quality control. Cold Spring Harb Perspect Biol. 2011;3(8):a004374.

24. Bukau B, Weissman J, Horwich A. Molecular chaperones and protein quality control. Cell. 2006;125(3):443–51.

25. Horianopoulos LC, Kronstad JW. Chaperone Networks in Fungal Pathogens of Humans. J Fungi (Basel). 2021;7(3).

26. Cowen LE, Lindquist S. Hsp90 potentiates the rapid evolution of new traits: drug resistance in diverse fungi. Science. 2005;309(5744):2185–9.

27. Padro-Villegas L, Mora-Montes HM. Heat Shock Proteins in Medically Relevant Fungal Pathogens: From Molecular Chaperones to Virulence Factors and Therapeutic Targets. J Fungi (Basel). 2026;12(6).

28. Yakubu UM, Morano KA. Roles of the nucleotide exchange factor and chaperone Hsp110 in cellular proteostasis and diseases of protein misfolding. Biol Chem. 2018;399(10):1215–21.

29. Shaner L, Morano KA. All in the family: atypical Hsp70 chaperones are conserved modulators of Hsp70 activity. Cell Stress Chaperones. 2007;12(1):1–8.

30. Zuo D, Subjeck J, Wang XY. Unfolding the Role of Large Heat Shock Proteins: New Insights and Therapeutic Implications. Front Immunol. 2016;7:75.

31. Easton DP, Kaneko Y, Subjeck JR. The hsp110 and Grp1 70 stress proteins: newly recognized relatives of the Hsp70s. Cell Stress Chaperones. 2000;5(4):276–90.

32. Liu Q, Hendrickson WA. Insights into Hsp70 chaperone activity from a crystal structure of the yeast Hsp110 Sse1. Cell. 2007;131(1):106–20.

33. Albanese V, Yam AY, Baughman J, Parnot C, Frydman J. Systems analyses reveal two chaperone networks with distinct functions in eukaryotic cells. Cell. 2006;124(1):75–88.

34. Shorter J. The mammalian disaggregase machinery: Hsp110 synergizes with Hsp70 and Hsp40 to catalyze protein disaggregation and reactivation in a cell-free system. PLoS One. 2011;6(10):e26319.

35. Yam AY, Albanese V, Lin HT, Frydman J. Hsp110 cooperates with different cytosolic HSP70 systems in a pathway for de novo folding. J Biol Chem. 2005;280(50):41252–61.

36. Yakubu UM, Morano KA. Suppression of aggregate and amyloid formation by a novel intrinsically disordered region in metazoan Hsp110 chaperones. J Biol Chem. 2021:100567.

37. Fan Q, Park KW, Du Z, Morano KA, Li L. The role of Sse1 in the de novo formation and variant determination of the [PSI+] prion. Genetics. 2007;177(3):1583–93.

38. Shaner L, Wegele H, Buchner J, Morano KA. The yeast Hsp110 Sse1 functionally interacts with the Hsp70 chaperones Ssa and Ssb. J Biol Chem. 2005;280(50):41262–9.

39. Liu XD, Morano KA, Thiele DJ. The yeast Hsp110 family member, Sse1, is an Hsp90 cochaperone. J Biol Chem. 1999;274(38):26654–60.

40. Goeckeler JL, Stephens A, Lee P, Caplan AJ, Brodsky JL. Overexpression of yeast Hsp110 homolog Sse1p suppresses ydj1-151 thermosensitivity and restores Hsp90-dependent activity. Mol Biol Cell. 2002;13(8):2760–70.

41. Hrizo SL, Gusarova V, Habiel DM, Goeckeler JL, Fisher EA, Brodsky JL. The Hsp110 molecular chaperone stabilizes apolipoprotein B from endoplasmic reticulum-associated degradation (ERAD). J Biol Chem. 2007;282(45):32665–75.

42. Dragovic Z, Broadley SA, Shomura Y, Bracher A, Hartl FU. Molecular chaperones of the Hsp110 family act as nucleotide exchange factors of Hsp70s. EMBO J. 2006;25(11):2519–28.

43. Facciponte JG, Wang XY, Subjeck JR. Hsp110 and Grp170, members of the Hsp70 superfamily, bind to scavenger receptor-A and scavenger receptor expressed by endothelial cells-I. Eur J Immunol. 2007;37(8):2268–79.

44. Raviol H, Sadlish H, Rodriguez F, Mayer MP, Bukau B. Chaperone network in the yeast cytosol: Hsp110 is revealed as an Hsp70 nucleotide exchange factor. EMBO J. 2006;25(11):2510–8.

45. Mandal AK, Gibney PA, Nillegoda NB, Theodoraki MA, Caplan AJ, Morano KA. Hsp110 chaperones control client fate determination in the hsp70-Hsp90 chaperone system. Mol Biol Cell. 2010;21(9):1439–48.

46. Ravindran MS, Bagchi P, Inoue T, Tsai B. A Non-enveloped Virus Hijacks Host Disaggregation Machinery to Translocate across the Endoplasmic Reticulum Membrane. PLoS Pathog. 2015;11(8):e1005086.

47. Gao X, Carroni M, Nussbaum-Krammer C, Mogk A, Nillegoda NB, Szlachcic A, et al. Human Hsp70 Disaggregase Reverses Parkinson’s-Linked alpha-Synuclein Amyloid Fibrils. Mol Cell. 2015;59(5):781–93.

48. Torrente MP, Shorter J. The metazoan protein disaggregase and amyloid depolymerase system: Hsp110, Hsp70, Hsp40, and small heat shock proteins. Prion. 2013;7(6):457–63.

49. Rampelt H, Kirstein-Miles J, Nillegoda NB, Chi K, Scholz SR, Morimoto RI, et al. Metazoan Hsp70 machines use Hsp110 to power protein disaggregation. EMBO J. 2012;31(21):4221–35.

50. Mattoo RU, Sharma SK, Priya S, Finka A, Goloubinoff P. Hsp110 is a bona fide chaperone using ATP to unfold stable misfolded polypeptides and reciprocally collaborate with Hsp70 to solubilize protein aggregates. J Biol Chem. 2013;288(29):21399–411.

51. Kandasamy G, Andreasson C. Hsp70-Hsp110 chaperones deliver ubiquitin-dependent and -independent substrates to the 26S proteasome for proteolysis in yeast. J Cell Sci. 2018;131(6).

52. Moran C, Kinsella GK, Zhang ZR, Perrett S, Jones GW. Mutational analysis of Sse1 (Hsp110) suggests an integral role for this chaperone in yeast prion propagation in vivo. G3 (Bethesda). 2013;3(8):1409–18.

53. O’Driscoll J, Clare D, Saibil H. Prion aggregate structure in yeast cells is determined by the Hsp104-Hsp110 disaggregase machinery. J Cell Biol. 2015;211(1):145–58.

54. Nagao J, Cho T, Uno J, Ueno K, Imayoshi R, Nakayama H, et al. Candida albicans Msi3p, a homolog of the Saccharomyces cerevisiae Sse1p of the Hsp70 family, is involved in cell growth and fluconazole tolerance. FEMS Yeast Res. 2012;12(6):728–37.

55. Cho T, Toyoda M, Sudoh M, Nakashima Y, Calderone RA, Kaminishi H. Isolation and sequencing of the Candida albicans MSI3, a putative novel member of the HSP70 family. Yeast. 2003;20(2):149–56.

56. Wang Y, Li H, Sun C, Liu Q, Zhou L, Liu Q. Purification and biochemical characterization of Msi3, an essential Hsp110 molecular chaperone in Candida albicans. Cell Stress Chaperones. 2021;26(4):695–704.

57. Lee-Yoon D, Easton D, Murawski M, Burd R, Subjeck JR. Identification of a major subfamily of large hsp70-like proteins through the cloning of the mammalian 110-kDa heat shock protein. J Biol Chem. 1995;270(26):15725–33.

58. Oh HJ, Chen X, Subjeck JR. Hsp110 protects heat-denatured proteins and confers cellular thermoresistance. J Biol Chem. 1997;272(50):31636–40.

59. Taguchi YV, Gorenberg EL, Nagy M, Thrasher D, Fenton WA, Volpicelli-Daley L, et al. Hsp110 mitigates alpha-synuclein pathology in vivo. Proc Natl Acad Sci U S A. 2019;116(48):24310–6.

60. Hendrickson WA, Liu Q. Exchange we can believe in. Structure. 2008;16(8):1153–5.

61. Bracher A, Verghese J. The nucleotide exchange factors of Hsp70 molecular chaperones. Front Mol Biosci. 2015;2:10.

62. Andreasson C, Fiaux J, Rampelt H, Druffel-Augustin S, Bukau B. Insights into the structural dynamics of the Hsp110-Hsp70 interaction reveal the mechanism for nucleotide exchange activity. Proc Natl Acad Sci U S A. 2008;105(43):16519–24.

63. Polier S, Dragovic Z, Hartl FU, Bracher A. Structural basis for the cooperation of Hsp70 and Hsp110 chaperones in protein folding. Cell. 2008;133(6):1068–79.

64. Schuermann JP, Jiang J, Cuellar J, Llorca O, Wang L, Gimenez LE, et al. Structure of the Hsp110:Hsc70 nucleotide exchange machine. Mol Cell. 2008;31(2):232–43.

65. Xu X, Sarbeng EB, Vorvis C, Kumar DP, Zhou L, Liu Q. The unique peptide substrate binding properties of 110 KDA heatshock protein (HSP110) determines its distinct chaperone activity. J Biol Chem. 2012;287(8):5661–72.

66. Hu L, Sun C, Kidd JM, Han J, Fang X, Li H, et al. A first-in-class inhibitor of Hsp110 molecular chaperones of pathogenic fungi. Nat Commun. 2023;14(1):2745.

67. Sun C, Li Y, Kidd JM, Han J, Ding L, May AE, et al. Characterization of a New Hsp110 Inhibitor as a Potential Antifungal. J Fungi (Basel). 2024;10(11).

68. Kmiecik SW, Mayer MP. Molecular mechanisms of heat shock factor 1 regulation. Trends Biochem Sci. 2022;47(3):218–34.

69. Li J, Labbadia J, Morimoto RI. Rethinking HSF1 in Stress, Development, and Organismal Health. Trends Cell Biol. 2017;27(12):895–905.

70. Gomez-Pastor R, Burchfiel ET, Thiele DJ. Regulation of heat shock transcription factors and their roles in physiology and disease. Nat Rev Mol Cell Biol. 2018;19(1):4–19.

71. Dai C, Whitesell L, Rogers AB, Lindquist S. Heat shock factor 1 is a powerful multifaceted modifier of carcinogenesis. Cell. 2007;130(6):1005–18.

72. Morimoto RI. The heat shock response: systems biology of proteotoxic stress in aging and disease. Cold Spring Harb Symp Quant Biol. 2011;76:91–9.

73. Akerfelt M, Morimoto RI, Sistonen L. Heat shock factors: integrators of cell stress, development and lifespan. Nat Rev Mol Cell Biol. 2010;11(8):545–55.

74. Mendillo ML, Santagata S, Koeva M, Bell GW, Hu R, Tamimi RM, et al. HSF1 drives a transcriptional program distinct from heat shock to support highly malignant human cancers. Cell. 2012;150(3):549–62.

75. Pincus D. Regulation of Hsf1 and the Heat Shock Response. Adv Exp Med Biol. 2020;1243:41–50.

76. Fujimoto M, Takii R, Nakai A. Regulation of HSF1 transcriptional complexes under proteotoxic stress. Bioessays. 2023;45(7):e2300036.

77. Dai C, Sampson SB. HSF1: Guardian of Proteostasis in Cancer. Trends Cell Biol. 2016;26(1):17–28.

78. Lindquist S. The heat-shock response. Annu Rev Biochem. 1986;55:1151–91.

79. Craig EA. The heat shock response. CRC Crit Rev Biochem. 1985;18(3):239–80.

80. Suo C, Gao Y, Ding C, Sun T. The function and regulation of heat shock transcription factor in Cryptococcus. Front Cell Infect Microbiol. 2023;13:1195968.

81. Veri AO, Robbins N, Cowen LE. Regulation of the heat shock transcription factor Hsf1 in fungi: implications for temperature-dependent virulence traits. FEMS Yeast Res. 2018;18(5).

82. Yang DH, Jung KW, Bang S, Lee JW, Song MH, Floyd-Averette A, et al. Rewiring of Signaling Networks Modulating Thermotolerance in the Human Pathogen Cryptococcus neoformans. Genetics. 2017;205(1):201–19.

83. Xiao X, Zuo X, Davis AA, McMillan DR, Curry BB, Richardson JA, et al. HSF1 is required for extra-embryonic development, postnatal growth and protection during inflammatory responses in mice. EMBO J. 1999;18(21):5943–52.

84. Dong B, Jaeger AM, Thiele DJ. Inhibiting Heat Shock Factor 1 in Cancer: A Unique Therapeutic Opportunity. Trends Pharmacol Sci. 2019;40(12):986–1005.

85. Dong B, Jaeger AM, Hughes PF, Loiselle DR, Hauck JS, Fu Y, et al. Targeting therapy-resistant prostate cancer via a direct inhibitor of the human heat shock transcription factor 1. Sci Transl Med. 2020;12(574).

86. Neudegger T, Verghese J, Hayer-Hartl M, Hartl FU, Bracher A. Structure of human heat-shock transcription factor 1 in complex with DNA. Nature Structural & Molecular Biology. 2016;23(2):140-+.

87. Kwon-Chung KJ, Fraser JA, Doering TL, Wang Z, Janbon G, Idnurm A, et al. Cryptococcus neoformans and Cryptococcus gattii, the etiologic agents of cryptococcosis. Cold Spring Harb Perspect Med. 2014;4(7):a019760.

88. Kozubowski L, Heitman J. Profiling a killer, the development of Cryptococcus neoformans. FEMS Microbiol Rev. 2012;36(1):78–94.

89. Odds FC, Brown AJ, Gow NA. Antifungal agents: mechanisms of action. Trends Microbiol. 2003;11(6):272–9.

90. Shor E, Chauhan N. A case for two-component signaling systems as antifungal drug targets. PLoS Pathog. 2015;11(2):e1004632.

91. Willmund F, del Alamo M, Pechmann S, Chen T, Albanese V, Dammer EB, et al. The cotranslational function of ribosome-associated Hsp70 in eukaryotic protein homeostasis. Cell. 2013;152(1-2):196–209.

92. Park HS, Chow EW, Fu C, Soderblom EJ, Moseley MA, Heitman J, et al. Calcineurin Targets Involved in Stress Survival and Fungal Virulence. PLoS Pathog. 2016;12(9):e1005873.

93. Fu C, Donadio N, Cardenas ME, Heitman J. Dissecting the Roles of the Calcineurin Pathway in Unisexual Reproduction, Stress Responses, and Virulence in Cryptococcus deneoformans. Genetics. 2018;208(2):639–53.

94. Halgren TA. Identifying and characterizing binding sites and assessing druggability. J Chem Inf Model. 2009;49(2):377–89.

95. Trott O, Olson AJ. AutoDock Vina: improving the speed and accuracy of docking with a new scoring function, efficient optimization, and multithreading. J Comput Chem. 2010;31(2):455–61.

96. Eberhardt J, Santos-Martins D, Tillack AF, Forli S. AutoDock Vina 1.2.0: New Docking Methods, Expanded Force Field, and Python Bindings. J Chem Inf Model. 2021;61(8):3891–8.

97. Salentin S, Schreiber S, Haupt VJ, Adasme MF, Schroeder M. PLIP: fully automated protein-ligand interaction profiler. Nucleic Acids Res. 2015;43(W1):W443–7.

98. Schake P, Bolz SN, Linnemann K, Schroeder M. PLIP 2025: introducing protein-protein interactions to the protein-ligand interaction profiler. Nucleic Acids Res. 2025;53(W1):W463-W5.

99. Bhattacharya S, Sobel JD, White TC. A Combination Fluorescence Assay Demonstrates Increased Efflux Pump Activity as a Resistance Mechanism in Azole-Resistant Vaginal Candida albicans Isolates. Antimicrob Agents Chemother. 2016;60(10):5858–66.

100. Koplin A, Preissler S, Ilina Y, Koch M, Scior A, Erhardt M, et al. A dual function for chaperones SSB-RAC and the NAC nascent polypeptide-associated complex on ribosomes. J Cell Biol. 2010;189(1):57–68.

101. Tyanova S, Cox J. Perseus: A Bioinformatics Platform for Integrative Analysis of Proteomics Data in Cancer Research. Methods Mol Biol. 2018;1711:133–48.

102. Ge SX, Jung D, Yao R. ShinyGO: a graphical gene-set enrichment tool for animals and plants. Bioinformatics. 2020;36(8):2628–9.

