## Supplemental Figures and Tables for "HSF1 inhibition potentiates the antifungal activity and proteome instability induced by the fungal Hsp110 inhibitor 2H"

### Figure S1

**A**

```
hHSF1       13  SNVPAFLTKLWTLVSDPDTDALICWSPSGNSFHVFDQGQFAKEVLPKYFKHNNMASFVRQ
cnHSF1      78  MKVPAFLNKLYTMVSDPEVDDLIYWGENGDSFFVPNAELFGRELLPRWFKHSNFSSFVRQ
CTA8        275 KTRPAFVMKIWSMVNDPANHEYIRWNDDGKTFQVFHREDFMKVILPKYFKHNNFASFVRQ
                 . ***: *::::*.**  .  * *. .*.:* * .   * : :**::***.*::*****

hHSF1           LNMYGFRKVVHIEQGGLVKP----E-RDDTEFQHPCFLRGQEQLLENIKRKVT     120
cnHSF1          LNMYGFHKVPHLQSGALKNE----TPIELWEFANPYFKRGQPQLLTKVTRKNN     186
CTA8            LNMYGWHKVQDVANGTLNQNSDKNGQDEIWQFENPNFIKDREDLLDKIVRNKS     387
                *****::** .: .* * :        :  :* :* * :.: :** :: *: .
```

**B**

```
Percent Identity  Matrix

1: hHSF1       100.00    51.85    44.44
2: cnHSF1       51.85   100.00    43.12
3: CTA8         44.44    43.12   100.00
```

**Figure S1. Conservation of the DNA-binding domains (DBDs) of HSF1 proteins.**
**(A)** Sequence alignment of the DBDs of HSF1 proteins. Human HSF1 (hHSF1), HSF1 from *C. neoformans* (cnHSF1), and HSF1 from *C. albicans* (CTA8) were aligned using Clustal2.1. The identical and conserved residues are labeled with * and :, respectively.
**(B)** Percent identity matrix of the DBDs among hHSF1, cnHSF1, and CTA8. The matrix was created by Clustal2.1.

#### Figure S2

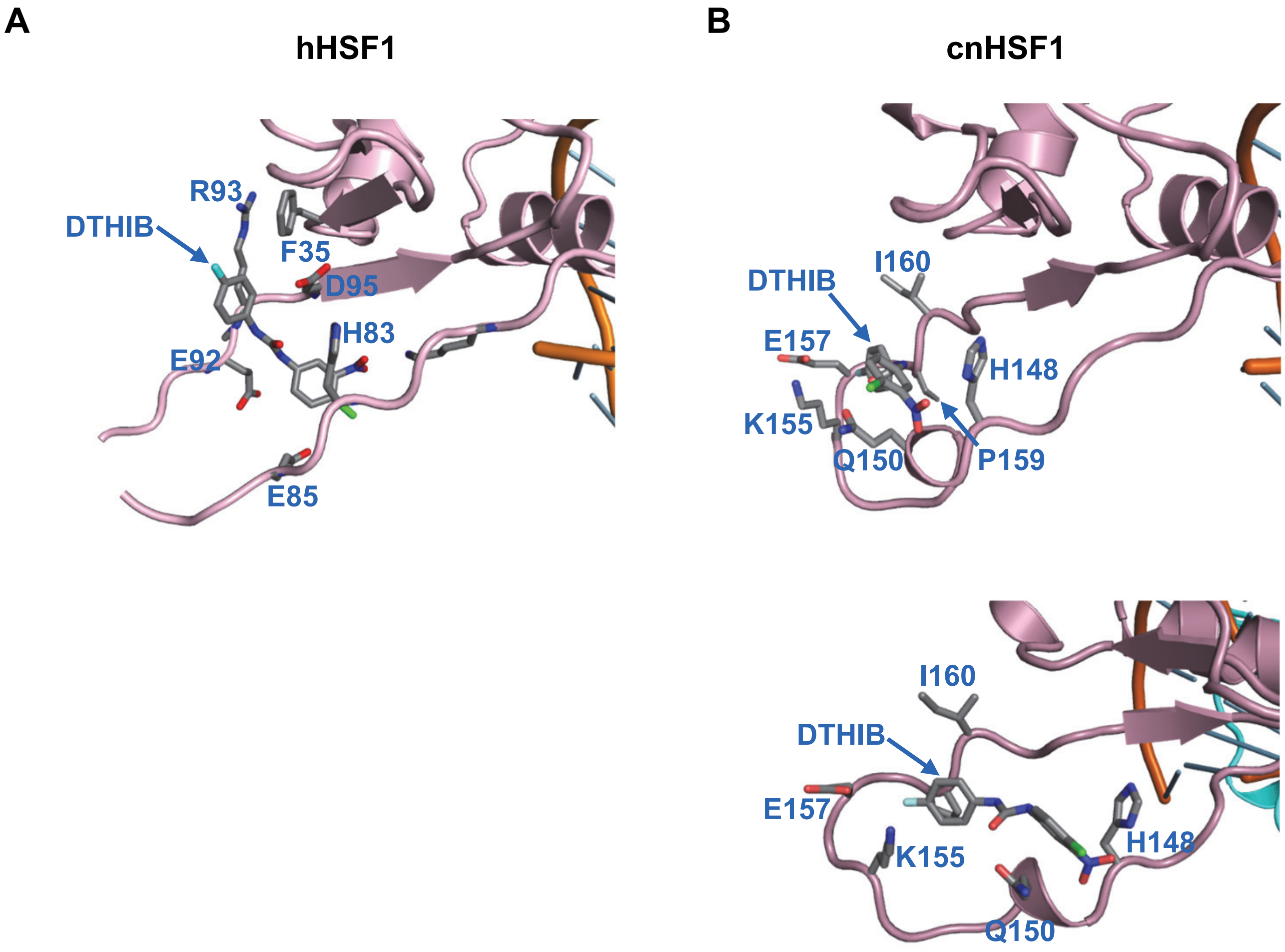


**Figure S2. Details of the molecular interactions between the DBDs of HSF1 proteins and docked DTHIB.**
**(A)** hHSF1; **(B)** cnHSF1 (two views). The cnHSF1 structure shown in the top panel of (**B**) is superimposed on the hHSF1 structure shown in (**A**). The cnHSF1 structure in the bottom panel of (**B**) is rotated to better visulalize the interacting residues. DTHIB and the contacting residues of the HSF1 proteins are shown in stick representation and labeled. PLIP analysis of the hHSF1-DTHIB complex identified a localized interaction environment involving F47, H83, E85, E92, R93, and D95. These residues provide chemically complementary interactions with DTHIB, including aromatic/hydrophobic, polar, acidic, and basic contributions. In the cnHSF1-DTHIB complex, the local interaction environment comprises H148, Q150, K155, E157, P159, and I160. These residues form a chemically diverse binding pocket around DTHIB. Thus, the cnHSF1 interaction pattern contains both polar and hydrophobic contributions and exhibits an overall physicochemical character similar to that of the hHSF1-DTHIB binding environment, although the individual interacting residues differ, with the exception of the conserved H83 in hHSF1 and the corresponding H148 in cnHSF1.

### Figure S3

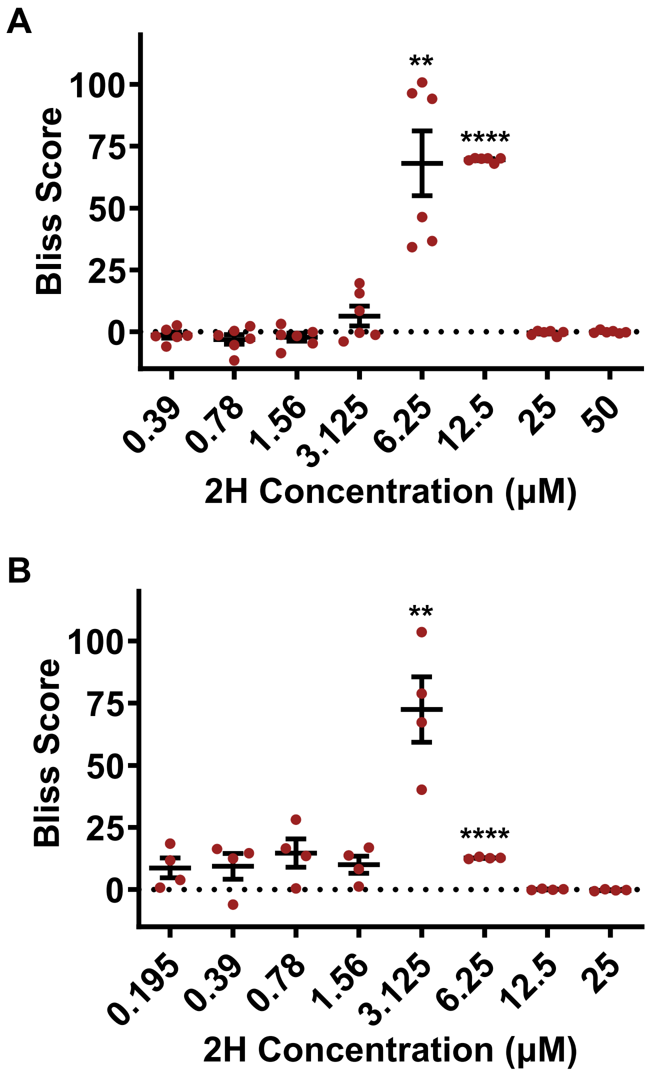


**Figure S3. Bliss synergy analysis of 2H combined with 25 μM DTHIB in *C. neoformans* (A) and *C. albicans* (B).**
Bliss scores were calculated from the growth inhibition data shown in **Fig. 2A** and **Fig. 2C** using the Bliss independence model. Relative growth values were normalized to vehicle-treated controls prior to analysis. Positive, zero, and negative Bliss scores indicate synergistic, additive, and antagonistic interactions, respectively. Each point represents an independent biological replicate (n = 6 for *C. neoformans* and n = 4 for *C. albicans*), and horizontal bars indicate mean ± SEM. Bliss scores were compared with the theoretical additive value of zero using a one-sample test, and P values were adjusted for multiple comparisons using the Benjamini-Hochberg false discovery rate (FDR) procedure. *P < 0.05; **P < 0.01; ****P < 0.0001. Detailed statistical results are provided in **Tables S4** and **S6**.

#### Figure S4

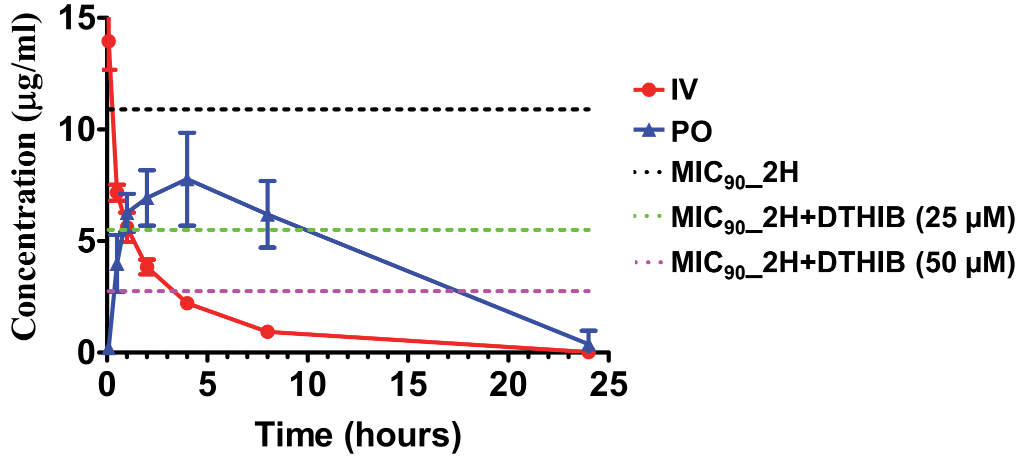


**Figure S4. Previously reported pharmacokinetic profile of 2H relative to antifungal activity thresholds.**

Plasma concentration-time profile of 2H following administration through intravenous injection (IV) and taken through the mouth (PO stands for per os). Institute of Cancer Research (ICR) mouse was used. The doses for IV and PO are 5 mg/kg and 20 mg/kg, respectively. The plots are presented as mean values ± SEM ( n=3 animals). Dashed lines indicate the $MIC_{90}$ values of 2H against *C. neoformans* H99 alone (black) and in combination with DTHIB (green, 25 μM; magenta, 50 μM). The reduction in $MIC_{90}$ produced by DTHIB increases the duration during which drug concentrations remain above the effective antifungal threshold.

### Figure S5

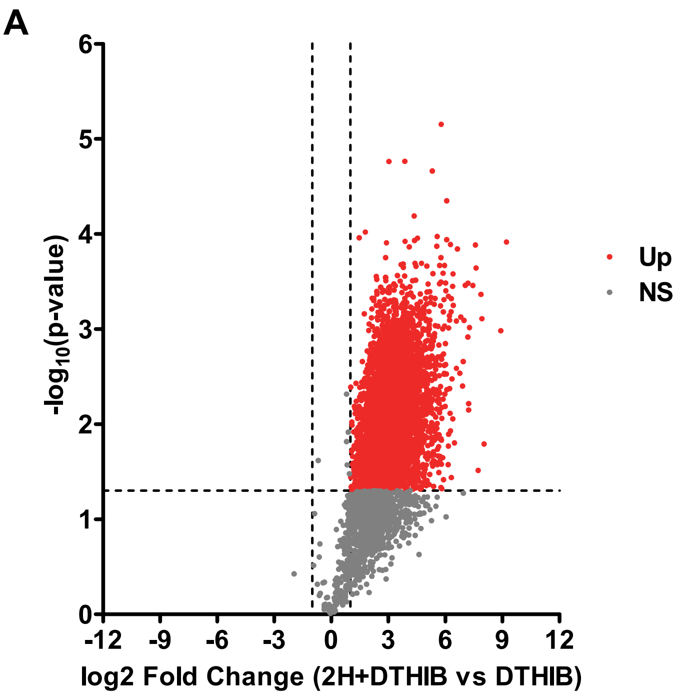


**Figure S5.** The volcano plot displays the $\log_2$ Fold Change in aggregate abundance for the combined treatment with 2H and DTHIB (2H+DTHIB) versus DTHIB treatment as a function of statistical significance ($-\log_{10}$ *P*-value). The volcano plot was analyzed in the same way as in **Fig. 5A**.

### Figure S6

**A**

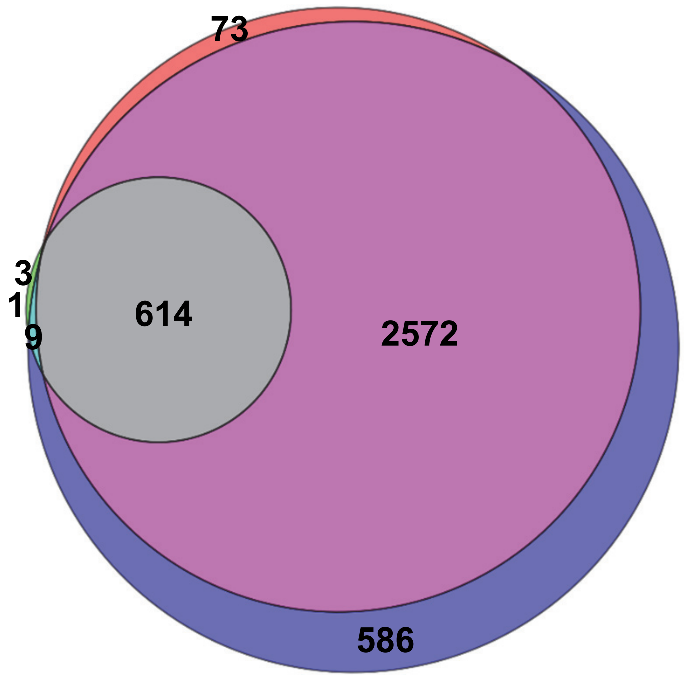


**B**

| | number |
|---|---|
| DTHIB only | 1 |
| 2H only | 73 |
| 2H+DTHIB only | 586 |
| DTHIB and 2H only | 3 |
| DTHIB and 2H+DTHIB only | 9 |
| 2H and 2H+DTHIB only | 2572 |
| DTHIB, 2H, and 2H+DTHIB common | 614 |

| Comparison | Overlapping proteins |
|---|---|
| DTHIB vs 2H | 617 |
| DTHIB vs 2H+DTHIB | 623 |
| 2H vs 2H+DTHIB | 3,186 |

**Figure S6. Comparison of the increased aggregated proteins (Up population) among the treatments.**

(**A**) The Venn diagram. 2H vs control (2H): red; combined treatment of 2H and DTHIB vs control (2H+DTHIB): blue; DTHIB vs control (DTHIB): green. The individual and overlapping numbers among the treatments are labeled.

(**B**) The list of the individual and overlapping numbers among the treatments for the Venn diagram in (**A**).

### Figure S7

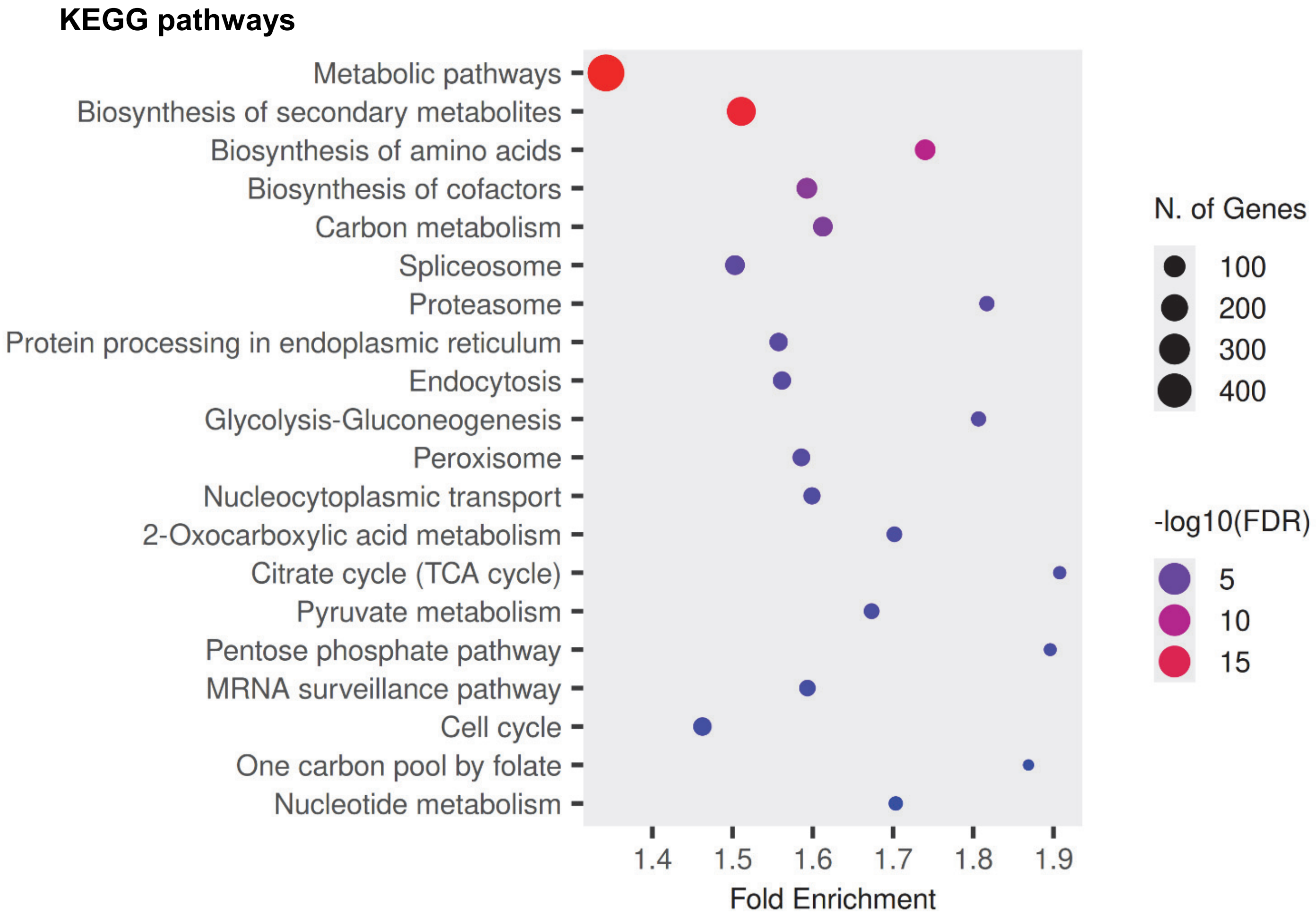


**Figure S7. The top 20 KEGG pathways enriched in the aggregated proteome of *C. neoformans* induced by 2H treatment.**
ShinyGO 8.6 (South Dakota State University) was used for KEGG pathway analysis and plot preparation.

### Figure S8

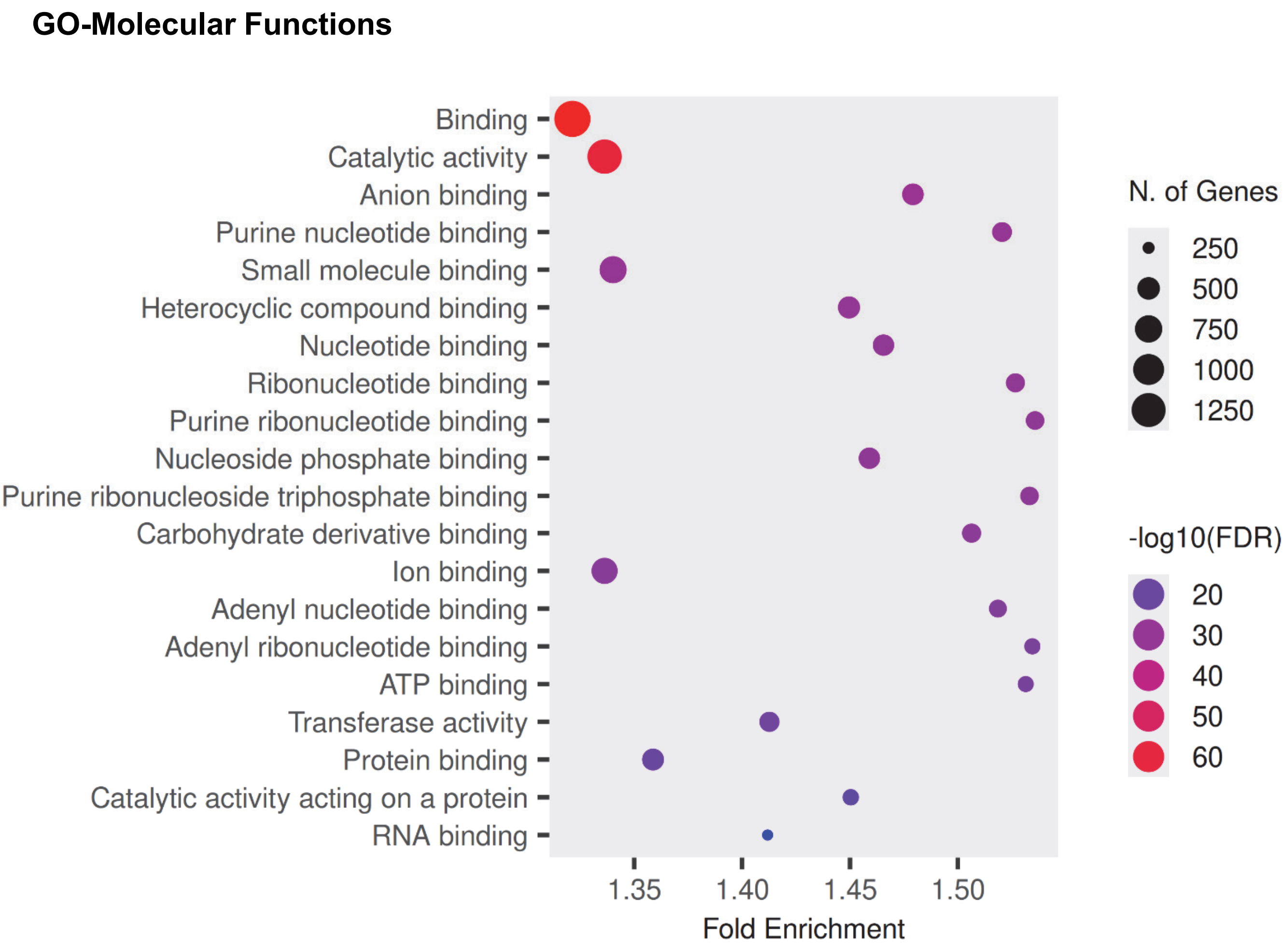


**Figure S8. The top 20 GO Molecular Functions enriched in the aggregated proteome of *C. neoformans* induced by 2H treatment.**
The plot was prepared using ShinyGO 8.6 (South Dakota State University).

### Table S1

| 2H (µM) | 2H vs. 2H + 25 µM DTHIB | Adjusted P value | 2H vs. 2H + 50 µM DTHIB | Adjusted P value |
|---|---|---|---|---|
| 0.39 | ns | 0.4396 | ns | 0.2337 |
| 0.78 | ns | 0.6213 | ns | 0.1042 |
| 1.56 | ns | 0.5471 | *** | 0.0005 |
| 3.125 | * | 0.0286 | **** | <0.0001 |
| 6.25 | **** | <0.0001 | **** | <0.0001 |
| 12.5 | **** | <0.0001 | **** | <0.0001 |
| 25 | ns | 0.9998 | ns | 0.9997 |
| 50 | ns | >0.9999 | ns | 0.9974 |

**Table S1. Sidak multiple-comparison analysis for Figure 2A.**
Statistical significance was determined by ordinary two-way ANOVA followed by Sidak's multiple-comparison test. Comparisons were performed between 2H alone and combination treatments at each concentration. ns, not significant; *P < 0.05; **P < 0.01; ***P < 0.001; ****P < 0.0001.

### Table S2

| 2H (µM) | 2H vs. 2H + 25 µM DTHIB | Adjusted P value | 2H vs. 2H + 50 µM DTHIB | Adjusted P value |
|---|---|---|---|---|
| 0.20 | ns | >0.9999 | ns | 0.9816 |
| 0.39 | ns | 0.9999 | ns | 0.9039 |
| 0.78 | ns | 0.8235 | ns | 0.7460 |
| 1.56 | ns | 0.9960 | *** | 0.0006 |
| 3.12 | **** | <0.0001 | **** | <0.0001 |
| 6.25 | ns | 0.2970 | ns | 0.2919 |
| 12.5 | ns | >0.9999 | ns | >0.9999 |
| 25 | ns | >0.9999 | ns | >0.9999 |

**Table S2. Sidak Multiple-Comparison Analysis of Figure 2C.**
Statistical significance was determined by ordinary two-way ANOVA followed by Sidak's multiple-comparison test. Comparisons were performed between 2H alone and combination treatments at each concentration. ns, not significant; *P < 0.05; **P < 0.01; ***P < 0.001; ****P < 0.0001.

### Table S3

| 2H (µM) | Mean Bliss Score | Raw P value | BH-FDR adjusted P value | Significant after FDR? |
|---|---|---|---|---|
| 0.39 | 8.31 | 0.0161 | 0.0215 | Yes |
| 0.78 | 10.69 | 0.0198 | 0.0238 | Yes |
| 1.56 | 21.72 | 0.0159 | 0.0215 | Yes |
| 3.125 | 34.68 | 0.0279 | 0.0319 | Yes |
| 6.25 | 103.7 | <0.0001 | <0.0004 | Yes |
| 12.5 | 75.12 | <0.0001 | <0.0004 | Yes |
| 25 | -0.55 | 0.1327 | 0.1327 | No |
| 50 | -1.11 | 0.0713 | 0.0815 | No |

**Table S3. Bliss synergy analysis for Figure 2B (2H + 50 µM DTHIB).**
Bliss scores were compared with the theoretical additive value of zero. Raw P values were obtained from one-sample tests against zero. Multiple-comparison correction was performed using the Benjamini-Hochberg false discovery rate procedure (FDR = 5%).

### Table S4

| 2H (µM) | Mean Bliss Score | Raw P value | BH-FDR adjusted P value | Significant after FDR? |
|---|---|---|---|---|
| 0.39 | -1.330 | 0.3143 | 0.3592 | No |
| 0.78 | -3.082 | 0.1830 | 0.2928 | No |
| 1.56 | -2.185 | 0.2434 | 0.3245 | No |
| 3.125 | 6.335 | 0.1728 | 0.2928 | No |
| 6.25 | 68.11 | 0.0035 | 0.0140 | Yes |
| 12.5 | 69.60 | <0.0001 | <0.0004 | Yes |
| 25 | -0.4813 | 0.2849 | 0.3592 | No |
| 50 | -0.0779 | 0.7566 | 0.7566 | No |

**Table S4. Bliss Synergy Analysis for Figure S3A (2H + 25 µM DTHIB).**
Bliss scores were compared with the theoretical additive value of zero. Raw P values were obtained from one-sample tests against zero. Multiple-comparison correction was performed using the Benjamini-Hochberg false discovery rate procedure (FDR = 5%).

### Table S5

| 2H (µM) | Mean Bliss Score | Raw P value | BH-FDR adjusted P value | Significant after FDR? |
|---|---|---|---|---|
| 0.2 | -1.854 | 0.6470 | 0.7394 | No |
| 0.4 | -3.831 | 0.4879 | 0.6505 | No |
| 0.8 | -5.945 | 0.4871 | 0.6505 | No |
| 1.6 | 28.80 | 0.0255 | 0.0510 | No* |
| 3.1 | 78.75 | 0.0029 | 0.0116 | Yes |
| 6.3 | 11.75 | <0.0001 | <0.0008 | Yes |
| 12.5 | -0.0665 | 0.7599 | 0.7599 | No |
| 25 | 0.0334 | 0.8103 | 0.8103 | No |

**Table S5. Bliss synergy analysis for Figure 2D (2H + 50 µM DTHIB).**
Bliss scores were compared with the theoretical additive value of zero. Raw P values were obtained from one-sample tests against zero. Multiple-comparison correction was performed using the Benjamini-Hochberg false discovery rate procedure (FDR = 5%).

### Table S6

| 2H (µM) | Mean Bliss Score | Raw P value | BH-FDR adjusted P value | Significant after FDR? |
|---|---|---|---|---|
| 0.195 | 8.717 | 0.1168 | 0.2336 | No |
| 0.39 | 9.349 | 0.1703 | 0.2479 | No |
| 0.78 | 14.69 | 0.0818 | 0.2181 | No |
| 1.56 | 10.02 | 0.0617 | 0.2181 | No |
| 3.125 | 72.49 | 0.0118 | 0.0472 | Yes |
| 6.25 | 12.68 | <0.0001 | <0.0008 | Yes |
| 12.5 | 0.0787 | 0.5765 | 0.6589 | No |
| 25 | -0.2796 | 0.1860 | 0.2479 | No |

**Table S6. Bliss Synergy Analysis for Figure S3B (2H + 25 µM DTHIB).**
Bliss scores were compared with the theoretical additive value of zero. Raw P values were obtained from one-sample tests against zero. Multiple-comparison correction was performed using the Benjamini-Hochberg false discovery rate procedure (FDR = 5%).

### Table S7

| Fluconazole (μM) | Mean Bliss Score | Raw P value | BH-FDR adjusted P value | Significant after FDR? |
|---|---|---|---|---|
| 0.5 | -0.776 | 0.8323 | 0.8323 | No |
| 1.0 | -9.173 | 0.0596 | 0.1073 | No |
| 2.0 | -8.200 | 0.0283 | 0.0637 | No |
| 3.9 | -20.21 | 0.0064 | 0.0192 | Yes |
| 7.8 | -76.22 | 0.0002 | 0.0018 | Yes |
| 15.6 | 0.584 | 0.5981 | 0.7177 | No |
| 31.25 | 0.705 | 0.1131 | 0.1697 | No |
| 62.5 | 0.010 | 0.9928 | 0.9928 | No |
| 125 | 0.578 | 0.0006 | 0.0027 | Yes |

**Table S7. Bliss Analysis for Figure 3B (Fluconazole + 50 μM DTHIB in *C. neoformans*).**
Bliss scores were compared with the theoretical additive value of zero. Raw P values were obtained from one-sample tests against zero. Multiple-comparison correction was performed using the Benjamini-Hochberg false discovery rate procedure (FDR = 5%).

### Table S8

| Fluconazole (µM) | Mean Bliss Score | Raw P value | BH-FDR adjusted P value | Significant after FDR? |
|---|---|---|---|---|
| 0.03125 | 0.656 | 0.8215 | 0.8215 | No |
| 0.0625 | 13.98 | 0.0203 | 0.0305 | Yes |
| 0.125 | -54.69 | 0.0003 | 0.0009 | Yes |
| 0.25 | -84.29 | <0.0001 | <0.0009 | Yes |
| 0.5 | -8.649 | 0.0092 | 0.0138 | Yes |
| 1 | 1.569 | 0.0298 | 0.0383 | Yes |
| 2 | 3.788 | 0.0002 | 0.0009 | Yes |
| 3.9 | 2.815 | 0.0003 | 0.0009 | Yes |
| 7.8 | 0.578 | 0.0006 | 0.0014 | Yes |

**Table S8. Bliss Analysis for Figure 3D (Fluconazole + 50 µM DTHIB in *C. albicans*).**
Bliss scores were compared with the theoretical additive value of zero. Raw P values were obtained from one-sample tests against zero. Multiple-comparison correction was performed using the Benjamini-Hochberg false discovery rate procedure (FDR = 5%).

### Table S9

| Treatment | Proteins detected |
|---|---|
| control | 2518 |
| | 3375 |
| | 3270 |
| 2H | 4495 |
| | 4569 |
| | 4515 |
| 2H+DTHIB | 4604 |
| | 4599 |
| | 4554 |
| DTHIB | 3575 |
| | 4074 |
| | 4031 |

**Table S9. Numbers of proteins detected in detergent-insoluble aggregates in three independent biological replicates.**
2H+DTHIB: combined treatment with 2H and DTHIB.

### Table S10

| 2H+DTHIB vs 2H | Mean (Da) | Median (50$^{th}$ percentile) (Da) | 25$^{th}$ percentile (Da) | 75$^{th}$ percentile (Da) |
|---|---|---|---|---|
| NS | 57,928 | 46,532 | 29,932 | 75,482 |
| Up | 67,496 | 60,893 | 40,021 | 88,962 |

**Table S10. Combined treatment with DTHIB further increases the molecular weight of the aggregated proteins induced by 2H treatment.**
Molecular weights for mean, median, 25$^{th}$ and 75$^{th}$ percentile are shown for the NS and Up populations of the aggregated proteins induced by 2H+DTHIB combined treatment versus the 2H treatment identified in the volcano plot in **Figure 6D**.

### Table S11

| Gene ID | Protein Function | Virulence Pathway | $log_2$ FC 2H | $log_2$ FC 2H+DTHIB |
|---|---|---|---|---|
| PTP2 | Protein tyrosine phosphatase | Stress-response signaling | 5.22 | 6.23 |
| CNAG_07338 | N-acyl-phosphatidylethanolamine-hydrolyzing phospholipase D | Stress-response signaling | 4.53 | 3.24 |
| CNAG_03786 | Lysophospholipase | Stress-response signaling | 4.41 | 5.51 |
| CNAG_00274 | Palmitoyltransferase | Stress-response signaling | 2.00 | 1.98 |
| CNAG_06782 | Alpha-1,2-mannosyltransferase | Cell wall and capsule biosynthesis | 3.53 | 4.48 |
| CNAG_00373 | Glucan 1,3-beta-glucosidase | Cell wall and capsule biosynthesis | 3.28 | 3.33 |
| CNAG_03832 | Alpha-1,2-mannosyltransferase | Cell wall and capsule biosynthesis | 3.06 | 3.55 |
| CNAG_00456 | Adhesin domain-containing protein | Cell wall and capsule biosynthesis | NC | 3.41 |
| CNAG_00742 | Serine C-palmitoyltransferase | Sphingolipid biosynthesis/metabolism | 4.89 | 4.77 |
| CNAG_00101 | Inositolphosphorylceramide synthase subunit Kei1-domain-containing protein | Sphingolipid biosynthesis/metabolism | 2.80 | 4.17 |
| CNAG_00644 | Sphingolipid 4-desaturase | Sphingolipid biosynthesis/metabolism | 1.62 | 4.61 |
| CNAG_00253 | Exocyst complex component SEC15 | Vesicle trafficking / exocytosis | 4.34 | 6.50 |
| CNAG_06792 | Protein-vacuolar targeting-related protein | Vesicle trafficking / exocytosis | 3.14 | 4.02 |
| CNAG_06738 | Exocyst complex protein EXO70 | Vesicle trafficking / exocytosis | 2.96 | 5.76 |
| CNAG_00363 | Histidine kinase | Two-component histidine kinase signaling | 4.00 | 7.36 |
| CNAG_00106 | Histidine kinase | Two-component histidine kinase signaling | 1.73 | 4.71 |
| CNAG_00802 | Calcineurin-binding protein | Calcineurin signaling | 3.82 | 5.37 |
| CNAG_00135 | Cyclophilin | Calcineurin signaling | 2.13 | 5.29 |
| CNAG_00570 | cAMP-dependent protein kinase regulatory subunit | cAMP/PKA signaling | 3.33 | NC |
| CNAG_00792 | ATP-binding cassette (ABC) transporter | Drug resistance transport systems | 3.04 | 3.87 |

**Table S11. Enriched virulence-associated proteins in *C. neoformans* following 2H treatment and combined treatment with 2H and DTHIB.**
Proteins were selected based on known or strongly supported roles in cryptococcal virulence. $log_2$ Fold Change ($log_2$ FC) values are listed for 2H versus control and for the combined treatment of 2H and DTHIB versus control (2H+DTHIB). NC indicates no significant change.
